# Unbiased screen of human transcriptome reveals an unexpected role of 3′UTRs in translation initiation

**DOI:** 10.1101/2021.05.28.446092

**Authors:** Yun Yang, Xiaojuan Fan, Yanwen Ye, Chuyun Chen, Sebastian E.J. Ludwig, Sirui Zhang, Qianyun Lu, Cindy L. Will, Henning Urlaub, Jing Sun, Reinhard Lührmann, Zefeng Wang

## Abstract

Although most eukaryotic mRNAs require a 5′-cap for translation initiation, some can also be translated through a poorly studied cap-independent pathway. Here we developed a circRNA-based system and unbiasedly identified more than 10,000 sequences in the human transcriptome that contain Cap-independent Translation Initiators (CiTIs). Surprisingly, most of the identified CiTIs are located in 3′UTRs, which mainly promote translation initiation in mRNAs bearing highly structured 5′UTR. Mechanistically, CiTI recruits several translation initiation factors including eIF3 and DHX29, which in turn unwind 5′UTR structures and facilitate ribosome scanning. Functionally, we showed that the translation of HIF1A mRNA, an endogenous DHX29 target, is antagonistically regulated by its 5′UTR structure and a new 3′-CiTI in response to hypoxia. Therefore, deletion of 3′-CiTI suppresses cell growth in hypoxia and tumor progression *in vivo*. Collectively, our study uncovers a new regulatory mode for translation where the 3′UTR actively participate in the translation initiation.

## Introduction

The mRNA translation is an essential and energetically expensive process tightly regulated at the initiation stage. In eukaryotic cells, canonical translation initiation is mediated by the formation of eIF4F complex on the 5′-cap (7-methylguanosine), followed by the attachment of the ribosomal 43S preinitiation complex that scans along the mRNA in a 5′-to-3′ direction for a start codon ^1,2^. Multiple *cis-*elements and *trans-*factors are employed to accurately regulate mRNA translation initiation. For example, the RNA structures and upstream ORF (uORF) in 5′UTRs are well known *cis*-elements that regulate mRNA translation, whereas many protein factors or miRNAs can control translation *in trans* ^1-4^.

In addition, many genes can also be translated in a cap-independent fashion, mostly through direct recruitment of ribosomes to the internal ribosome entry site (IRES), which often occurs during stress conditions (like virus infection, heat shock, or apoptosis) ^5,6^. It has been estimated that up to 10-15% of cellular mRNAs undergo cap-independent translation under various conditions where cap-dependent translation is suppressed ^7^. A recent high-throughput screen targeting the viral RNAs and 5′UTRs of human mRNAs revealed that many human genes (583 out of 5058 genes tested) harbor the “cap-independent translation elements” that function as IRESs to drive mRNA translation ^8^. However, an unbiased screen of the translation initiation elements remains absent. Recently, thousands of circular RNAs (circRNAs) have been identified in eukaryotic cells ^9-11^, many of which can be translated *in vitro* and *in vivo* ^12-16^. Since circRNAs lack free ends and can only be translated cap-independently, they may serve as an improved system to study this noncanonical translation pathway.

## Results

### Identification of cap-independent translation initiation elements with a circRNA-based translation system

To unbiasedly screen the human transcriptome for endogenous sequences that drive cap-independent translation, we developed an efficient cell-based circRNA translation system (Fig. 1a) that has been extensively validated ^13,15-17^. We inserted a library of cDNA short fragments from normalized human transcriptome before start codon of the “split” EGFP (Fig. 1a) ^13,16^, and transfected the library to generate ∼3 million stable clones. The green cells were selected by FACS (Extended data Fig. 1a), and the insertion libraries were sequenced at each stage (supplementary information).

**Figure 1.**
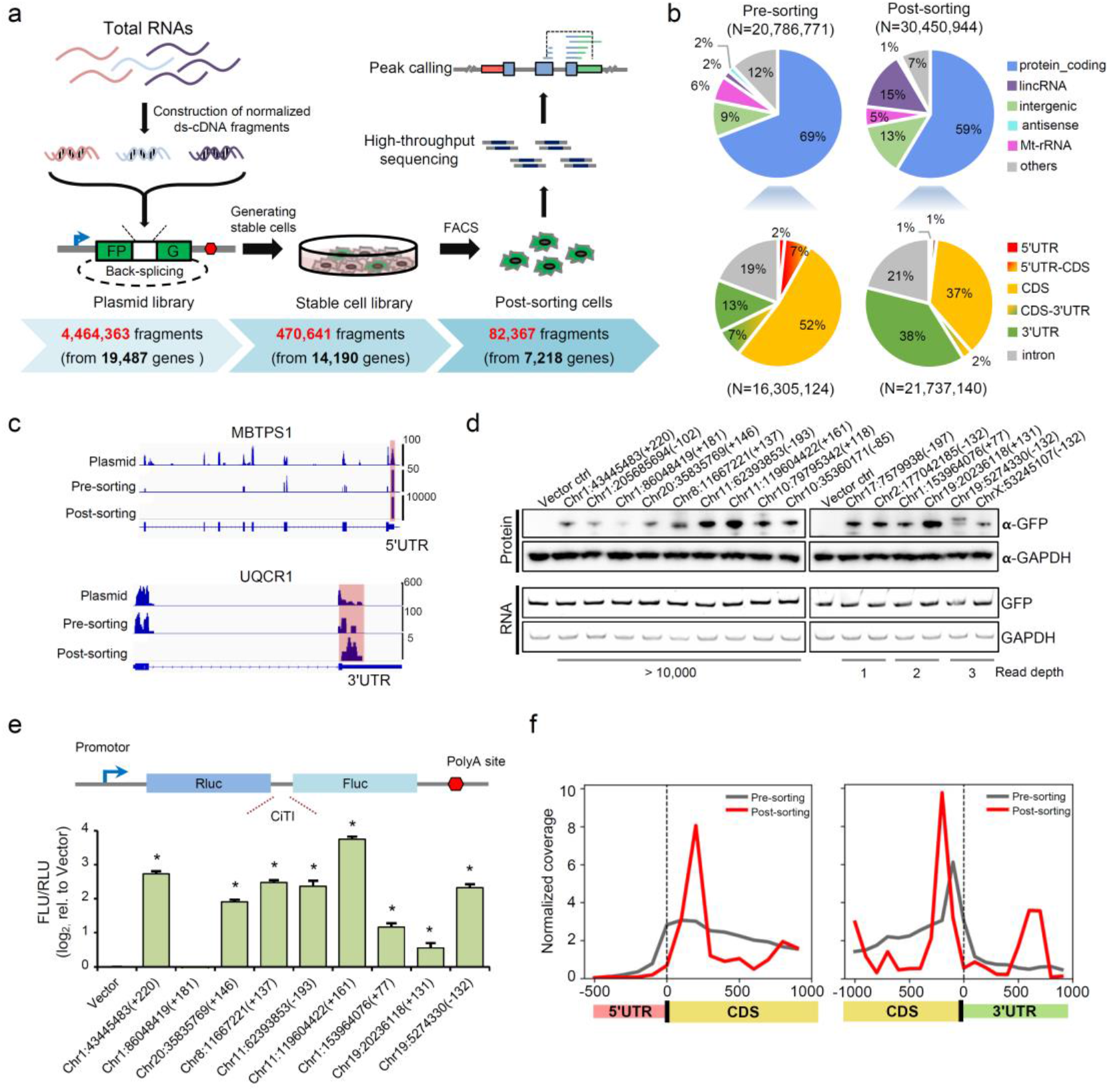
Systematic identification of cap-independent translation elements. **a**, Unbiased screen of cap-independent translation sequences using a circRNA reporter containing a split EGFP. Total HeLa RNAs were fragmented into 200-300nt fragments, reverse-transcribed, normalized and inserted into the circRNA reporter with FlpIn backbone to generate a plasmid library. This library was co-transfected with pOG44 (ratio 1:9) into Flp-In™-293 cells and selected by hygromycin to generate a library of stably transfected cells that were further sorted by FACS. The library was sequenced at each stage to determine sequence complexity. **b**, Genomic distribution of inserted sequences in pre-sorting and post-sorting cells (N indicates the number of fragments). **c**, The read coverage plots for MBTPS1 and UQCR10 as representative examples. **d**, Validation of the newly identified fragments (indicated by starting position in human genome and offset) using circRNA translation reporters. The fragments of both high (>10,000) and low (1 to 3) read depth were selected for testing. **e**, Validation of the newly identified fragments for cap-independent translation activation using bicistronic translation reporters inserted with different CiTIs. The reporters were transfected into HEK293 cells, and the products were determined by luminescence assay (n = 3, mean ± SD, * indicated p<0.05 by Student’s t-test). **f**, Positional density plots of the pre-sorting fragments and the CiTIs were in different regions of the gene body.

The fragment abundance in this library correlated poorly with gene levels in HeLa cells (r<0.09, Extended data Fig. 1b), with rRNAs representing <10% of the total fragments (Extended data Fig. 1c), indicative of an effective transcriptome normalization. After sorting, we recovered 82,367 unique fragments from the green cells with similar length distribution as the input (Extended data Fig. 1d and Extended data Table 1). The recovered fragments were not dominated by the pre-sorting library (Extended data Fig. 1e), suggesting the screen was unbiased. The majority of the reads from the pre- and post-sorting libraries were located in protein coding genes (Fig. 1b), with two examples shown in read density plots (Fig. 1c). Surprisingly, the percentage of positive reads from 3′UTRs was substantially increased whereas the reads from 5′UTRs and coding sequences (CDS) was decreased after sorting (Fig. 1b), suggesting a prominent role of 3′UTRs in regulation of cap-independent translation.

We further validated 14 out of 15 randomly selected fragments for their activity in driving circRNA translation (Fig. 1d, activities are independent on their read depth), indicating a low false positive rate. The IRES-like activities in 8 out of 9 positive fragments were also confirmed using a bicistronic translation reporter (Fig. 1e). Conversely, 3 out of the randomly selected 12 single clones from the input library promoted circRNA translation (Extended data Fig. 1f), suggesting that our screen had not reached saturation.

The overlapping positive fragments were merged into 10,455 peaks in human transcriptome, which were defined as cap-independent translation initiators (CiTIs). Most CiTIs were recovered only by a small number of reads in the post-sorting library while several CiTIs have exceedingly high read depth (Extended data Fig. 1g). The CiTIs were widely distributed across all human chromosomes (Extended data Fig. 1h), and their abundance correlated positively with the gene numbers (r=0.96) and the size (r=0.76) of each chromosome (Extended data Fig. 1i). The newly identified CiTIs included many previously reported IRESs ^8,18^ (Extended data Fig. 1j), including ∼10% of IRESs identified from a screen biased towards the 5′UTRs and viral sequences ^8^.

The CiTI-containing genes were more conserved than an average human gene (Extended data Fig. 2a), and were actively expressed and translated in cultured cells (Extended data Fig. 2b). Interestingly, CiTI containing genes were substantially enriched in functional categories of RNA translation, splicing, cell-cell adhesion and viral processes (Extended data Fig. 2c). Particularly, the CiTIs were enriched in genes encoding ribosomal proteins and translation initiation factors, suggesting a potential positive feedback regulation of translation.

Although the canonical IRESs are mostly located in the 5′UTR, the newly identified CiTIs were found in all gene regions including 5′UTRs (termed 5′-CiTI), CDS (termed C-CiTI), and 3′UTRs (termed 3′-CiTI). Consistent with our earlier finding, the CiTIs were enriched in 3′UTRs and peaked in the CDS near the start and stop codons (Fig. 1f). Compared to the cognate controls, we found a significant increase of sequence conservation in 3′-CiTIs, a modest increase in 5′-CiTIs, and no increase in C-CiTIs (Extended data Fig. 2d). The CiTIs generally had less structures as judged by higher minimal free energy (MFE) (Extended data Fig. 2e), whereas the canonical IRESs were significantly more structured ^18^, suggesting that, unlike IRESs, most CiTIs do not function by forming a specific structure.

Several enriched motifs from the CiTIs, including the previously reported U-rich motifs ^8,19^, were identified and further experimentally validated (Extended data Fig. 3a-3d). Consistent with previous reports ^8,20^, we found that the 5′-CiTIs tend to interact with 18S rRNAs *via* base-paring, and such interaction is sufficient to drive cap-independent translation (Extended data Fig. 3e-3g). Therefore, our circRNA-based translation system successfully identified several categories of CiTIs that contain both previously confirmed elements/motifs and novel elements enriched within 3′UTRs.

### Genes with 3′-CiTIs tend to contain downstream ORFs or structured 5′UTRs

The surprising enrichment of CiTIs in 3′UTRs rather than 5′UTRs implied an unexpected new role of 3′UTR in promoting translation initiation. Naturally, the 3′-CiTI may function as a *bona fide* IRES to drive cap-independent translation of downstream ORFs (dORF). By analyzing all potential dORFs, we found that the dORFs in the genes with 3′-CiTIs were more conserved than those without 3′-CiTIs (Extended data Fig. 4a), suggesting that 3′-CiTIs are probably under functional selection to drive dORF translation. Further analysis using published datasets of ribosome footprints ^21,22^ showed that a significant fraction of these dORFs (177/564) also had an upstream 3′-CiTI (Extended data Fig. 4b, p=2.9×10^−29^). However, we only identified the potential translation products from 7 of these dORFs using public mass spectrometry datasets ^23,24^. The lack of strong proteomic evidences may be due to cell-or tissue-specific translation of these dORFs, and/or the rapid degradation of these dORF-encoded proteins just like 3′UTR-coded peptides from translation read-through ^25,26^. These observations suggest that driving cap-independent translation of dORFs might not be the function for most 3′-CiTIs.

The fact that >90% of 3′-CiTI-containing genes lack evidence for dORF translation (Extended data Fig. 4b) suggested unknown roles of 3′-CiTIs. Interestingly, the 3′-CiTI-containing genes were significantly more conserved and more structured in their 5′UTRs than those without 3′-CiTIs (Fig. 2a-2b). They were also significantly overlapped with the genes containing upstream ORFs (uORF) (Extended data Fig. 4c) ^27,28^. Since most 5′UTR structures and uORFs play inhibitory roles in regulating translation ^29^, we speculated that 3′-CiTIs may counteract these 5′UTR structures or uORFs to enhance the main ORF (mORF) translation.

**Figure 2.**
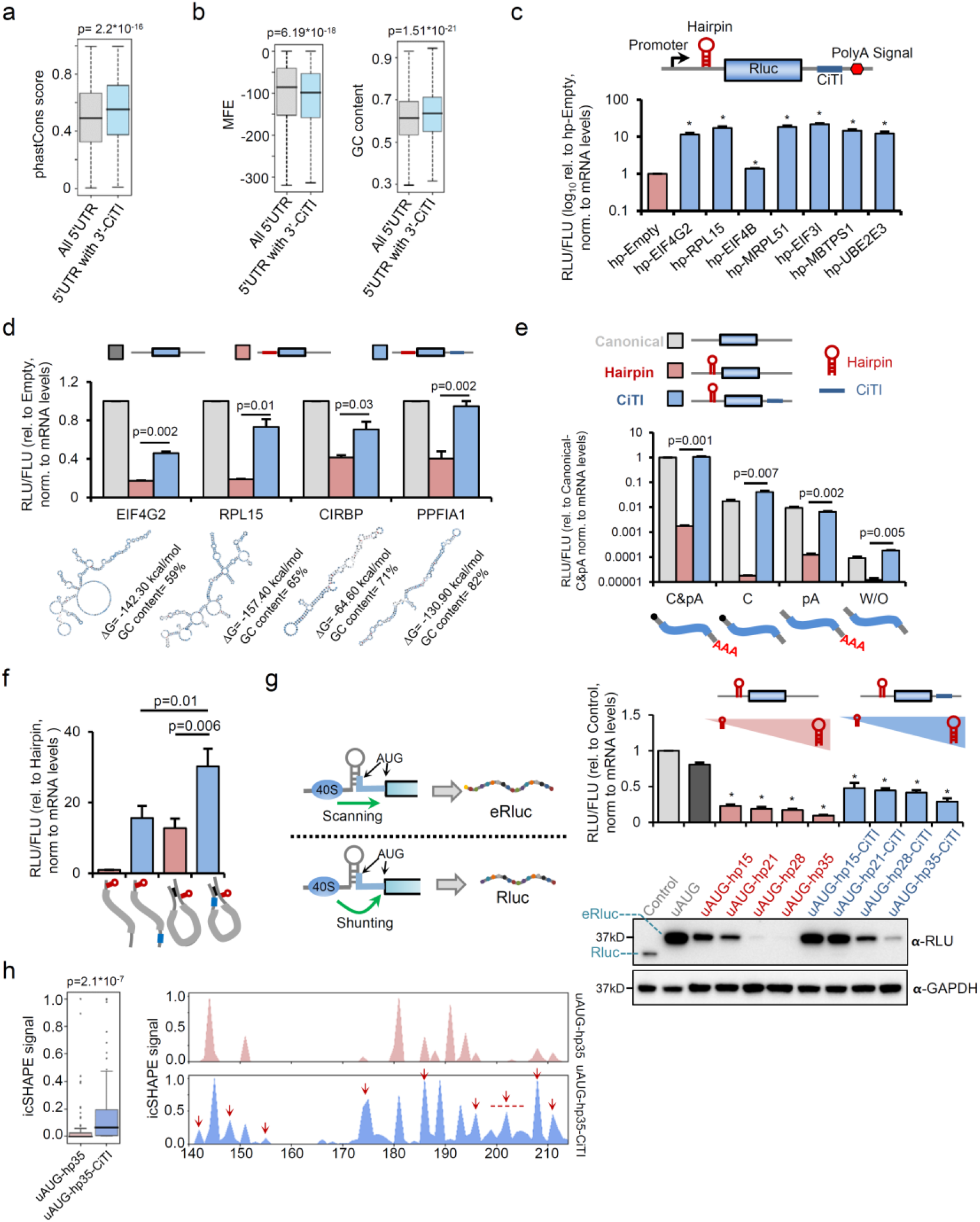
The 3′-CiTIs enhance translation of mRNAs with structured 5′UTRs. **a-b**, Sequence conservation the structure (predicted MFE, and GC contents) of the 5′UTRs from all human genes *vs*. the 3′-CiTI-containing genes. In **panels a**,**b and h**, p values were calculated by unpaired two-sided Mann–Whitney U test. **c**, The dual-luciferase reporter containing a hairpin at the 5′UTR of Rluc and different CiTIs in its 3′UTR. The translation efficiency of Rluc was determined by the relative abundance of proteins (*via* luminescence assay) *vs* RNAs (*via* qPCR) normalized to the control Fluc. In this and the following panels **(c-g)**, same procedures were used with Student’s t-test to evaluate statistical significance (n = 3, mean ± SD, * indicated p<0.05). **d**, Different combinations of endogenous structured 5′UTRs (shown as RNAfold prediction) and their cognate 3′-CiTIs were inserted into the Rluc reporter to test their effects on translation. **e**, Different Rluc mRNAs with an optional 5′-cap or poly-A were transcribed *in vitro*, and co-transfected with control Fluc mRNA into HEK293 cells to test translation efficiency. **f**, Introduction of mRNA looping through complementary sequences enhanced the activity of the 3′-CiTI. The experimental procedures are same as **panel e. g**, Two potential mechanisms for translation initiation through a 5′UTR structure (ribosome scanning *vs* shunting). The reporters with different structures in their 5′UTR and optional 3′-CiTIs were transfected into HEK293 cells, and the translation efficiency was determined by the relative abundance of Rluc proteins (*via* luminescence assay) vs RNAs (*via* qPCR) normalized to the control Fluc. **h**, *In vivo* icSHAPE signals of the hairpin regions in uAUG-hp35 mRNA with optional 3′-CiTI mRNAs. The icSHAPE signals of the hairpin region were plotted, and regions with increased opened structure by 3′-CiTI are indicated with red arrows.

Such possibility was directly tested using four translation reporters with different 5′UTRs (Extended data Fig. 5a): a canonical reporter with a regular 5′UTR, a hairpin reporter containing a 5′UTR hairpin immediately before the ORF, and two uORF reporters with the 5′UTRs containing two endogenous uORFs ^30,31^. We found that 6 out of 7 CiTIs inserted into the 3′UTR of the hairpin reporter significantly increased the translation (>10-fold increase) (Fig. 2c), and this translation enhancement was independent of the CiTI length or the cell lines used (Extended data Fig. 5b-5c). In contrast, no reliable translation increasement or even inhibitory effect was detected when CiTIs were inserted into the canonical reporter (Extended data Fig. 5d). As another comparison, insertion of CiTIs into the 3′UTR of the uORF reporters showed a weaker effect (Extended data Fig. 5e).

We next tested several pairs of highly structured 5′UTR and 3′-CiTI from endogenous mRNAs for their activities to cooperatively regulate translation. As expected, the structured 5′UTRs from all endogenous genes suppressed translation, and such suppression could be “reversed” by the cognate 3′-CiTIs (Fig. 2d), again suggesting a general translation enhancer activity of 3′-CiTIs that counteracts the translation suppression mediated by secondary structures in the 5′UTR.

Most human mRNAs contain a 5′-cap and poly-A tail that are looped together by a protein complex to increase translation efficiency ^32-34^. To determine if this looping is required for 3′-CiTI activity, we transfected cells with *in vitro* synthesized mRNAs containing optional 5′-cap or poly-A tail (Fig. 2e). In all mRNAs tested, insertion of a 3′-CiTI reversed the translation suppression by the 5′-hairpin, indicating the translation promotion by 3′-CiTI is independent of the 5′-cap or poly-A tail (Fig. 2e). This finding was further validated with a PolI reporter that can produce RNAs without a 5′-cap or poly-A tail, where a 3′-CiTI again enhanced translation of these “naked” mRNAs containing a 5′-hairpin (Extended data Fig. 6). Interestingly, the translation of mRNA that lacks a 5′-cap or poly-A tail could be enhanced by simply introducing a mRNA loop across the ORF using complementary sequences, and insertion of a 3′-CiTI further increased the translation of such mRNAs (Fig. 2f).

### 3′-CiTIs unwind the 5′UTR structures to facilitate ribosome scanning

During translation initiation, the preinitiation complex (PIC) usually needs to scan through the 5′UTR structures to initiate translation ^2,35-37^. Occasionally, the PIC can “jump over” the 5′UTR structures, a phenomenon called ribosome shunting ^38^ (Fig. 2g, left). To determine how 3′-CiTIs enhance translation of mRNAs containing 5′UTR structures, we generated several reporters with 5′-hairpins of different lengths and an upstream in-frame start codon (uAUG) inside the hairpin (Extended data Fig. 6a). The first start codon uAUG will be used by default to generate an extended Rluc (eRluc) *via* ribosome scanning. Alternatively, the PIC may “jump over” the structure and use the second AUG (i.e., without structure unwinding) to produce the canonical Rluc.

We found that the uAUG was exclusively used during translation, suggesting a ribosome scanning in all reporter mRNAs (Fig. 2g). As expected, introduction of a 5′-hairpin reduced the translation efficiency in a length-dependent fashion, and insertion of a 3′-CiTI increased the translation from the uAUG as judged by both western blots and luciferase activities (Fig. 2g). This increasement was achieved *via* promoting ribosome scanning rather than ribosome shunting, as only the proximal start codon was used in all cases. Additionally, we found that the reporter mRNA containing a 3′-CiTI had less structures (as judged by icSHAPE signals) than the mRNA without the 3′-CiTI, in both the entire 5′UTR and its hairpin region (Fig. 2h, and Extended data Fig. 7b-c), suggesting that the 3′-CiTI may unwind the upstream structures to help ribosomes scan through the 5′UTR.

### *Trans*-acting factors are involved in 3′-CiTI-mediated translation enhancement

To determine if the 3′-CiTIs recruit *trans*-acting factors to promote translation, we performed RNA affinity purification (Fig. 3a) using the MS2-fused RNAs containing three different 3′-CiTIs and controls (Fig. 2c, Extended data Fig. 8a). The purified RNP complexes were analyzed with mass spectrometry to identify 63 candidate proteins that specifically bind all three CiTIs (Extended data Fig. 8b and Table 2). These CiTI-binding proteins formed a densely-connected protein-protein interaction network with key functions in translation initiation and RNA processing (Fig. 3b), including many components in the eIF2 or eIF3 complex that functions as a multi-tasking machine to promote translation ^39^. Interestingly, the newly identified CiTI-binding proteins also include DHX29 (Fig. 3b), a DExH-box RNA helicase that might help to unwind RNA structures.

**Figure 3.**
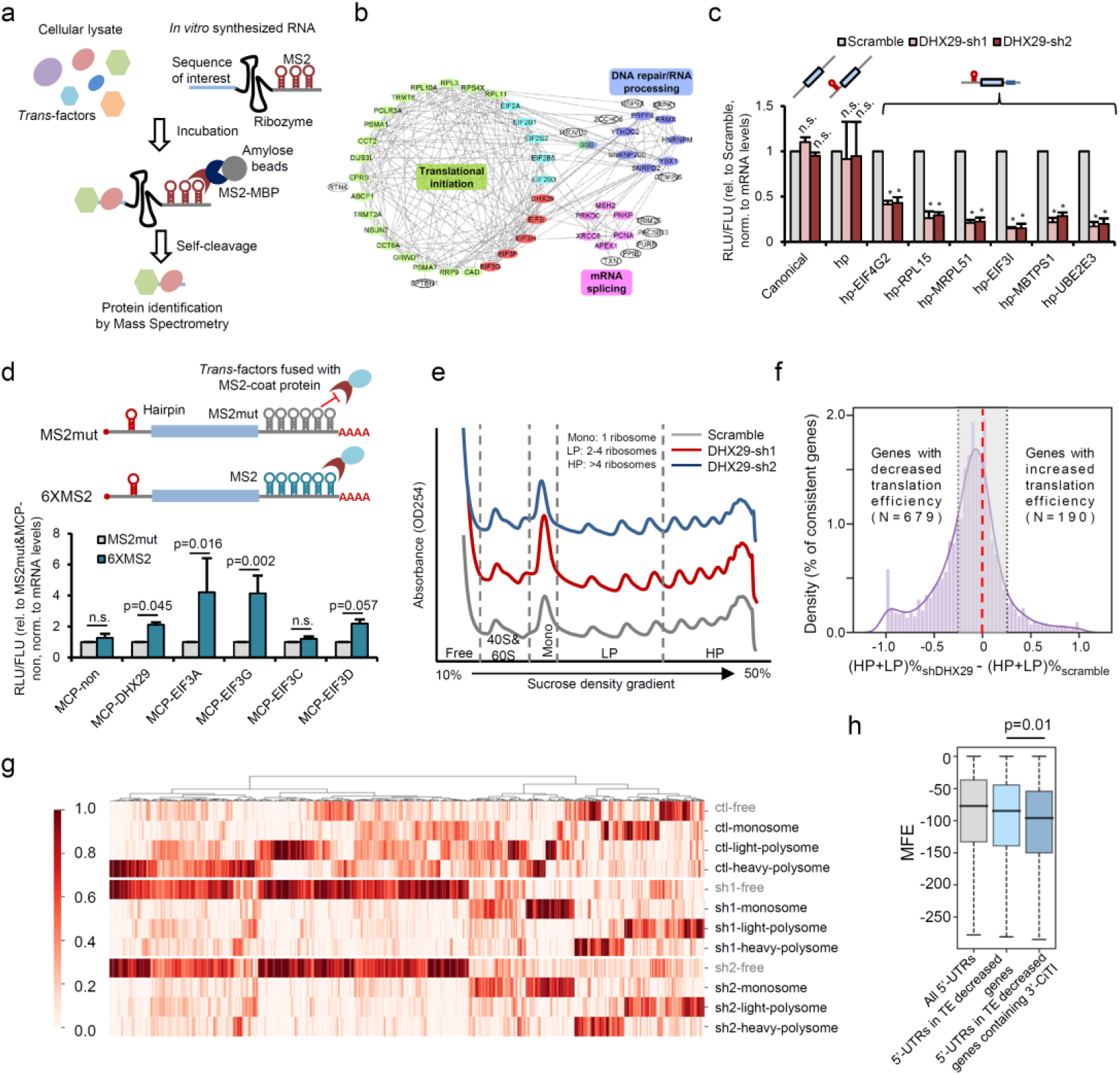
*Trans*-factors involved in 3′-CiTI functions. **a**, Schematic diagram of RNP affinity purification. **b**, The protein-protein interaction network of identified CiTI-binding *trans*-factors. **c**, DHX29 knockdown reduces the 3′-CiTI activity. The reporters with a 5′UTR hairpin and different 3′-CiTIs were transfected into HEK293 cells after DHX29 knockdown, and the translation efficiency of Rluc was determined as in Fig. 2c (n = 3, mean ± SD in **panel c and d**, * indicates p<0.05 by Student’s t-test, n.s. = not significant). **d**, Tethering of different *trans*-factors promotes translation of mRNAs with structured 5′UTRs. The translation reporter containing 6xMS2 sites was co-transfected with expression vectors of different fusion proteins, and the translation efficiency was measured as in **panel c. e**, Polysome profiling of SH-SY5Y cells after DHX29 knockdown. Mono: monosome; LP: light polysome (2-4 ribosomes); HP: heavy polysome (>4 ribosomes). The RNAs from each fraction were sequenced using the ribo-minus protocol. **f**, Global translation changes after DHX29-knockdown. The translation efficiency of each gene was defined by the percentage of polysome-associated mRNAs (data of **panel e**). The genes with consistent profiles in both knockdown lines were selected to plot percent difference of the polysome-associated mRNAs between knockdown cells *vs*. control cells (± 25% cutoffs were indicated). **g**, Relative distributions in polysome profiles were plotted as heatmap. The genes harboring altered distribution in DHX29 knockdown SH-SY5Y cells were shown. **h**, The predicted MFE of 5′UTRs from genes with reduced translation efficiency by DHX29 knockdown (p values calculated with unpaired two-sided Mann–Whitney U test).

In line with the possible role of DHX29 in CiTI-mediated translation regulation, knockdown of DHX29 specifically reduced the translation of several reporter mRNAs containing 3′-CiTIs (Fig. 3c, Extended data Fig. 8c). Furthermore, directly tethering DHX29 onto the 3′UTR significantly increased translation of a hairpin-containing mRNA (Fig. 3d, Extended data Fig. 8d), suggesting that DHX29 recruitment is sufficient for translation promotion. The recruitment of several eIF3 components also promoted the translation (Fig. 3d), supporting a previous finding that DHX29 cooperates with eIF3 during translation initiation ^29,40^. Since several eIF3 components were also pulled down by 3′-CiTIs (Fig. 3b), DHX29 may function together with the eIF3 complex to mediate the CiTI activity.

Previously, DHX29 was shown to regulate the translation of a handful of genes, including Cdc25C and XIAP ^41,42^. However, ∼2/3 of human mRNAs have structured 5′UTRs (i.e., with low MFE, ΔG < −40 kcal/mol) ^42^, many of which may also be regulated by DHX29. To systematically identify *bona fide* DHX29 targets, we performed polysome profiling after DHX29 knockdown, and sequenced mRNAs from different gradient fractions (Fig. 3e and Extended data Fig. 8c). We focused on the genes with consistent profiles in two independent DHX29 knockdown lines (Pearson correlation > 0.8), and calculated the translation efficiency by the percent of polysome-associated mRNAs ^43^. In total, 869 genes were identified with at least a 25% change in translation efficiency (Fig. 3f), most of which showed a translation reduction (Fig. 3f and 3g, Extended data Table 3). Furthermore, the genes with reduced translation efficiency in DHX29 knockdown cells had more RNA structures in their 5′UTR (Fig. 3h), and 91 of these genes also contained 3′-CiTIs (p=0.001 by hypergeometric test). Collectively, these results suggested that DHX29 plays a critical role in unwinding mRNA structures and promotes translation upon specific recruitment by 3′-CiTIs.

### 3′-CiTI and DHX29 promote HIF1A translation in response to hypoxia *in vitro* and *in vivo*

An important new target of DHX29 is the hypoxia-induced transcription factor HIF1A, which contains both a 3′-CiTI and strong 5′UTR structure ^44,45^. As a key factor for cellular response to hypoxia, HIF1A translation is suppressed in normoxia but strongly induced by oxygen deprivation ^46-48^. Using translation reporters containing HIF1A 5′UTR and different fragments of its 3′UTR, we found that the 5′UTR significantly suppressed translation under hypoxia, however the HIF1A 3′-CiTI fully restored translation in two different cell lines (Fig. 4a, top). Conversely, deletion of the 3′-CiTI from the HIF1A 3′UTR reduced translation under hypoxia (Fig. 4a, bottom). In contrast, under normoxia condition, translation enhancement by the 3′-CiTI was observed only in HCT116 cells (Extended data Fig. 9a-b).

**Figure 4.**
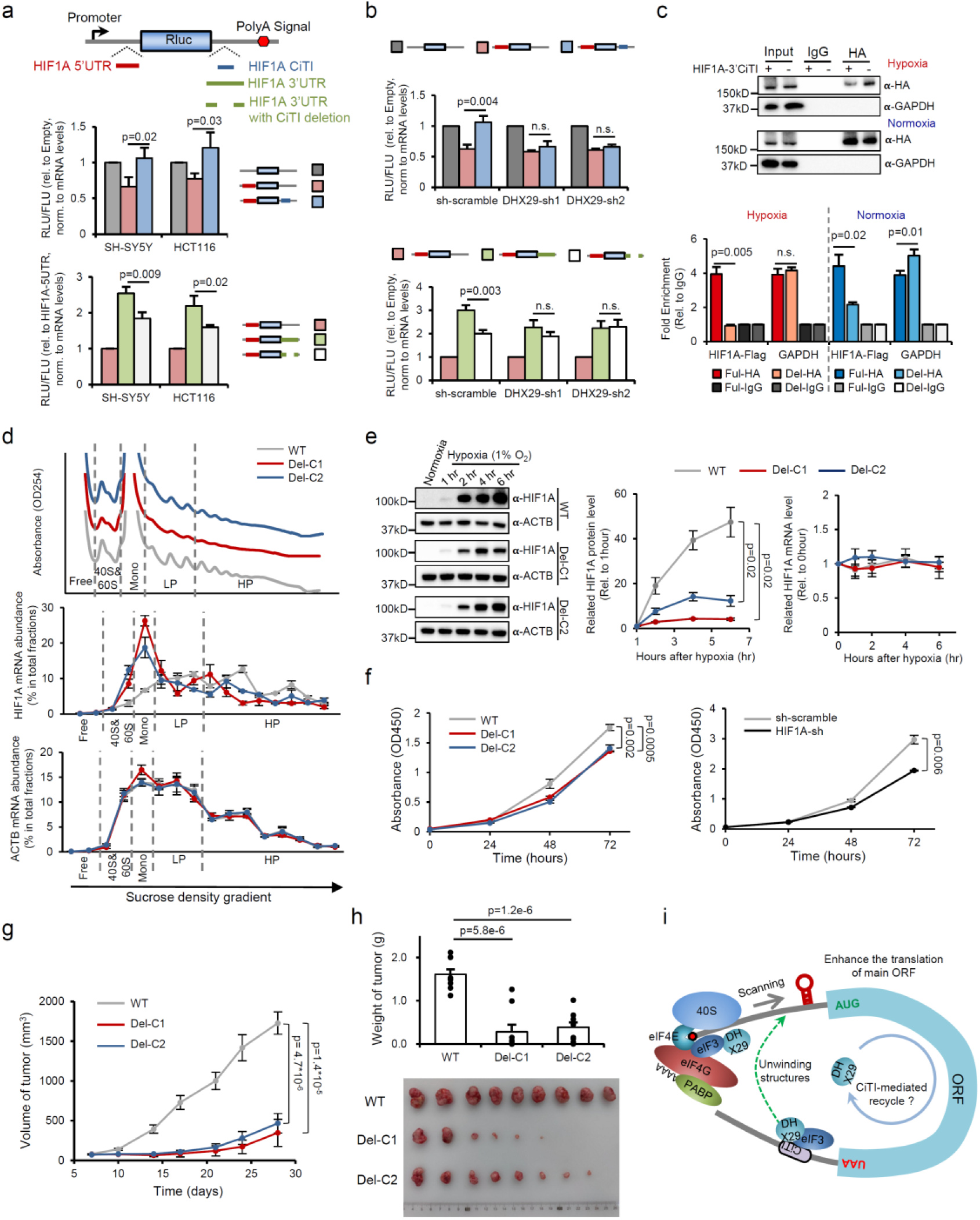
Biological function of HIF1A 3′-CiTI. **a**, Different translation reporters containing various combinations of the 5′UTR and 3′-CiTI from HIF1A were transfected in SH-SY5Y or HCT116 cells, and the translation efficiencies were examined as in Fig. 2c under hypoxia (1% O_2_) (n = 3, mean ± SD, Student’s t-test in **panel a-b**). **b**, The activity of the HIF1A 3′-CiTI was reduced in SH-SY5Y cells after DHX29-knockdown under hypoxia (1% O_2_). **c**, HA-tagged DHX29 was co-transfected with flag-tagged HIF1A (full length or 3′-CiTI deletion) into SH-SY5Y cells, and the interaction of HIF1A RNA and DHX29 was measured by RNA-IP and qPCR (n = 3, mean ± SD, Student’s t-test). **d**, The SH-SY5Y cells with deletion of HIF1A 3′-CITI were analyzed with polysome profiling after 6 h under hypoxia. Top, polysome profiling of WT and two independent deletion clones. Middle and bottom, the mRNA abundance of HIF1A and ACTB control in each fraction. **e**, WT cells and two 3′-CITI deletion clones were collected at 1 h, 2 h, 4 h, and 6 h after hypoxia, and the protein or RNA level of HIF1A was determined by western blot (left) or RT-qPCR (right). Middle, quantification of western blot (n = 3, mean ± SD, Student’s t-test). **f**, left, proliferation of WT SH-SY5Y cells and cells with HIF1A 3′-CiTI deletion under hypoxia as measured with CCK8 assay (n = 3, mean ± SD, Student’s t-test). The HIF1A knockdown cells were used as a control (right). **g**, Growth of xenograft tumor generated with WT SH-SY5Y cells or two clones of HIF1A 3′-CiTI deletion (n = 9, mean ± SEM, Student’s t-test). **h**, The weights and images of the xenograft tumors 4 weeks after injection. **i**, New model for 3′-CiTI-mediated translation regulation.

The translation promotion by the HIF1A 3′-CiTI was reduced by DHX29 knockdown under hypoxia (Fig. 4b). We further observed a direct interaction of DHX29 with HIF1A mRNAs (Extended data Fig. 9c), consistent with its general function as a subunit of the translation initiation complex ^40^. Interestingly, the interaction between DHX29 and HIF1A mRNA was nearly eliminated by the 3′-CiTI deletion under hypoxia, whereas this interaction was only reduced by ∼50% during normoxia (Fig. 4c), suggesting that the 3′-CiTI is essential for DHX29 recruitment to HIF1A mRNA during hypoxia.

To evaluate the functional impact of 3′-CiTI-mediated translation promotion, we deleted the 3′-CiTI from endogenous HIF1A gene in SH-SY5Y neuroblastoma cells with CRISPR-cas9 (Extended data Fig. 9d). Under normoxia, HIF1A mRNA was mostly found in monosomes or light polysomes (Extended data Fig. 9e), consistent with its low translation efficiency under this condition. Hypoxia induced the association of most HIF1A mRNAs with polysomes (Fig. 4d). However, deletion of the 3′-CiTI shifted HIF1A mRNAs (but not control mRNAs) from the polysome factions to the monosome fractions (Fig. 4d), suggesting that 3′-CiTI is required to specifically promote translation of HIF1A mRNA in response to hypoxia.

Furthermore, deletion of the 3′-CiTI significantly blunted the induction of HIF1A protein by hypoxia (Fig. 4e), resulting in a small but significant reduction in cell proliferation similar to that in HIF1A knockdown cells (Fig. 4f and Extended data Fig. 9f). Importantly, when inoculated in nude mice, 3′-CiTI deleted SH-SY5Y neuroblastoma tumors exhibited a dramatic reduction of growth (Fig. 4g and 4h). Taken together, these observations indicate that the 3′-CiTI in HIF1A mRNA functions as a key *cis*-element that recruits DHX29 to mediate environmental regulation of its translation, which plays a critical role in supporting *in vivo* tumor growth.

## Discussion

By unbiasedly screening human transcriptome, we uncovered a new class of 3′UTR elements that play an unexpected role in translation initiation. Specifically, the 3′-CiTIs actively recruit key initiation factors to unwind inhibitory RNA structures in the 5′UTRs, thereby promoting the ribosomes scanning (Fig. 4i). Since actively translated mRNAs generally form a loop through interactions of poly-A binding proteins with eIF4G ^49^, the 3′UTR may generally serve as a repository of regulatory elements that “catch and store” the translation initiation factors to increase their local concentration for the next round of translation.

This new model not only reveals a new role of 3′UTRs in translation initiation, but also explains why the 3′-CiTIs were initially identified as cap-independent translation initiator in our circRNA screening system. Unlike the canonical translation initiated with formation of eIF4F complex on 5′-cap, the 3′-CiTI in circRNAs may recruit initiation factors (e.g., DHX29 and eIF3 and eIF2 components), thus hijacking the initiation machine to bypass the requirement of eIF4F and drive cap-independent translation. Consequently, these 3′-CiTIs are conceptually equivalent to the “IRES” when inserted into the suitable context, although they function as translation initiators/enhancers from 3′UTR in endogenous settings.

Our findings further indicate that the 3′-CiTI-mediated translational control is an ancient and common mode of regulation. Firstly, the 3′-CiTIs are significantly more conserved than the matched control regions (Extended data Fig. 1d) and the 5′UTR in the 3′-CiTI-containing mRNAs are also more conserved and more structured (Fig. 2a-2b), indicating that the pairs of 3′-CiTIs and 5′UTRs are under functional selection in evolution. Moreover, ∼44% of the 2681 newly identified 3′-CiTIs are located in human mRNAs with structured 5′UTRs (ΔG < −40 kcal/mol)^42^, suggesting a common role of 3′-CiTIs in counteracting the inhibitory 5′UTR structures in many genes.

This study also highlights the importance of DHX29 in 3′-CiTI-mediated translation initiation in response to environmental stress. Several RNA helicases (including DHX29, DHX9 and DDX3) were reported to promote translation ^29,40^. In particular, DHX29 has been shown to interact with eIF3 to form an active 43S/DHX29 complex ^41^. However, it is unclear how these helicases are specifically recruited to mRNAs. Our data indicate that many 3′-CiTIs can function as the *cis-*acting elements to recruit DHX29 (and eIF3). The biological impact of such regulation is exemplified by HIF1A: deletion of a 3′-CiTI on the endogenous *HIF1A* locus directly impairs its translation and reduces cell survival under hypoxia, which further blunts *in vivo* tumor progression.

Given the various stress conditions where the cap-dependent translation initiation is inhibited (e.g., hypoxia, viral infection, heat/cold shock, etc.), the cooperative regulation of translation by the structured 5′UTR and 3′-CiTI reported here adds an additional layer of regulatory complexity during stress. Future studies are needed to determine how CiTI-mediated translational initiation can interplay with other modes of translation control from 3′UTRs, such as regulation by m6A and miRNAs ^4,50^.

In summary, our study uncovers an uncanonical mechanism of 3′UTR in mediating translation initiation, and further reveals the importance of this regulation in stress-induced cell survival and proliferation *in vitro* and *in vivo*. These findings may pave the way for novel cancer therapies that target stress-induced translation.

## Data availability

The raw sequencing data and filtered datasets are publicly available from NODE (assess number: OEP001285) and NCBI (assess number: PRJNA680213). The customized scripts for data analyses are also available from GitHub.

## Acknowledgments

The authors thank Dr. Hao Chi for his support in identifying dORF peptides from proteomic data using open pFind(v3.1.3), Dr. Eran Segal and Dr. Shira Weingarten-Gabbay for kindly sharing the raw data of their previous screen, Dr. Qiangfeng Cliff Zhang for the suggestion in icSHAPE analysis, and Drs. Xiaoling Li, Dieter A. Wolf, Shu-Bing Qian and Nahum Sonenberg for useful discussions. This work is supported by the Natural Science Foundation of China to Z.W. (91940303, 31661143031, and 31730110) and Y.Y. (91753135, 31870814), the National Key Research and Development Program of China to Z.W. (2018YFA0107602), and funding from the Max Plank Society to H.U. and R.L. Z.W. is also supported by the type A CAS Pioneer 100-Talent program. Y.Y. is sponsored by the Youth Innovation Promotion Association CAS (2019267), SA-SIBS Scholarship Program and Shanghai Science and Technology Committee Rising-Star Program (19QA1410500).

