## Supplemental materials for "Unbiased screen of human transcriptome reveals an unexpected role of 3′UTRs in translation initiation"

### Summary of supplemental materials

#### Supplementary information

#### Experimental procedures

#### Extended data figures and tables

**Extended data figure 1**. The screen for human, endogenous *cis*-elements involved in cap-independent translation.
**Extended data figure 2**. Distinct features of human endogenous CiTIs.
**Extended data figure 3**. Short motifs of human endogenous CiTIs.
**Extended data figure 4**. Analysis of downstream ORFs in the 3′UTR and features in the 5′UTR of 3′-CiTI containing genes.
**Extended data figure 5**. The diverse activities of 3′-CiTIs in different reporter systems.
**Extended data figure 6**. Validation of CiTI activity using PolI reporter.
**Extended data figure 7**. The 3′-CiTIs help to unwind 5′UTR structures.
**Extended data figure 8**. Identification of *trans*-factors binding to CiTIs.
**Extended data figure 9**. The function of the 3′-CiTI in HIF1A mRNAs under normal conditions.

**Extended data Table 1**. Newly identified CiTIs, related to Figure 1. (see attached excel file)
**Extended data Table 2**. *Trans*-factors binding to CiTIs identified by Mass Spectrometry, related to Figure 5. (see attached excel file)
**Extended data Table 3**. Genes with differential translation efficiency in DHX29 knockdown cells, related to Figure 5. (see attached excel file)
**Extended data Table 4**. Primers and sequences used in this study, related to Methods section (see attached excel file)

#### Supplementary information

##### Construction and screen of normalized transcriptome library for sequences that drive cap-independent circRNA translation.

IRES activity was mostly examined with bicistronic reporters that would introduce false discoveries derived from cryptic promoter or splicing sites. Although various controls can be used to reduce false discoveries [1], the existence of cellular IRESs was often challenged and the scope of cap-independent translation in the human transcriptome is still unclear. Recently, thousands of circular RNAs (circRNAs), most generated through pre-mRNA back-splicing, have been identified in eukaryotic cells [2-4]. Intriguingly, many circRNAs can be translated cap-independently (due to lack of free ends). For instance, the circRNAs containing a viral IRES were found to be translated *in vitro* [5] and inside cells [6], and many endogenous circRNAs were reported to be translated through cellular IRESs [7,8] or N(6)-methyladenosine modified (m6A) sites [9]. Since circRNAs lack free ends, they could serve as an improved system to study cap-independent translation.

To unbiasedly screen the human transcriptome for endogenous sequences that drive cap-independent translation, we developed a cell-based circRNA translation system (Fig. 1a, and Methods) that has been extensively validated by multiple groups [6,8-10]. We constructed a library of cDNA short fragments from a normalized transcriptome of HeLa cells, and inserted the library before the start codon of a GFP ORF in the circRNA reporter [6,9]. The resulting library was transfected into

Flp-In™-293 cells to generate ~3 million stable clones with a single reporter inserted in each clone (see Methods). The cells with strong GFP signals were collected by fluorescence-activated cell sorting (FACS), and the insertion libraries were sequenced before and after cell sorting (Extended data Fig. 1a).

The starting library contained 4,464,363 short unique fragments from 19,487 genes, with an average length around 200 nt (Fig. 1a, Extended data Fig.1d, and Extended data Table 1). The fragment abundance in this library correlated poorly with gene expression levels in HeLa cells ($r<0.09$, Extended data Fig. S1b), and the rRNA fragments represented less than 10% of the total library (Extended data Fig. 1c), suggesting that the transcriptome was effectively normalized during library construction. From the stable transfected cells, we recovered 82,367 unique fragments (~18%) from the green cells with similar length distribution (Extended data Fig.1d, and Extended data Table 1). The rRNA fragments were further reduced to less than 0.5% of total reads upon sorting (Extended data Fig. 1c). The majority of the reads from the pre- and post-sorting libraries were located in protein coding genes (Fig.1b). Intriguingly, the percentage of positive reads from 3′UTRs was substantially increased whereas the percentage of reads from 5′UTRs and coding sequences (CDS) was decreased in sorted strong GFP positive cells (Fig. 1b).

**The enriched motifs from the newly identified CiTIs**

Using a statistic enrichment analysis [11,12], we identified several motifs enriched in the newly identified CiTIs from different mRNA regions, including 5′-CiTIs, C-CiTIs and 3′-CiTIs (Extended data Fig. 3a). Among them, the enrichment of U-rich and C/U rich motifs in IRESs was also reported previously [13,14], and these motifs may function by directly recruiting the polypyrimidine track-binding proteins [15]. We tested this possibility by inserting poly(U)$_{10}$ into cap-independent translation reporters and co-expressing them with several RNA binding proteins (RBPs) that are known to specifically bind pyrimidine-rich sequences (PTBP1, HuR, RALY, hnRNPCL1) [16-18]. As expected, PTBP1 showed a robust activity to promote cap-independent translation in both circRNA or bicistronic reporters (Extended data Fig. 3b-c).

To control for the different binding affinity of these RBPs to the poly-U motif, we directly tethered these RBPs using the Puf domain specifically bound to an 8-nt sequence in the bicistronic mRNA (Extended data Fig. 3d), and again found that PTBP1 robustly promoted translation, with an activity comparable to that of the direct tethering of a ribosomal protein (Extended data Fig. 3d). Interestingly, direct tethering of HuR and hnRNPCL1 can also enhance cap-independent activity in this system (Extended data Fig. 3d). Together, these observations exemplify a general regulatory mechanism for cap-independent translation, in which the RBPs may promote translation by directly binding to specific motifs (such as poly-U or C/U motifs) in mRNA and function as *trans*-activators.

#### Many 5′-CITIs drive cap-independent translation by pairing to 18S rRNA

Previously, several IRESs were found to drive cap-independent translation by base-pairing with 18S rRNA [13,19-24]. To examine if the 5′-CiTIs function through a similar mechanism, we compared human 18S rRNA and the CiTIs for complementary sequences, and found that the 5′-CiTIs, but not 3′-CiTIs or C-CiTIs, are enriched for sequences that can base-pair with 18S rRNA (Extended data Fig. 3e). This result is consistent with an earlier report that sequences complementary to 18S rRNA are enriched in the 5′UTRs of human and yeast genes [25].

Among the three 18S rRNA regions that pair with 5′-CiTIs, two regions (position 650 and 1100) covered helix 18 and helix 26 that were previously reported to interact with mRNAs to promote cap-independent translation [13,26-29]. To experimentally validate the activity of these sequences, we selected three 5′-CiTIs that are paired with each of the three 18S rRNA regions, and tested their activities using two cap-independent translation reporters. As a positive control, we also included a 5′-CiTI containing a well-known short IRES (Gtx) that is paired to 18S rRNA [13,19-24]. All CiTIs tested can promote cap-independent translation in a circRNA or bicistronic reporter, and such activities were largely abolished by mutations disrupting their complementarity to 18S rRNA (Extended data Fig. 3f-g). This result suggests that the 18S rRNA complementary sequences in the 5′-CiTIs can function as IRESs, which is consistent with previous findings (Malygin et al., 2013; Matsuda and Mauro, 2014; Chappell et al., 2000;Weingarten-Gabbay et al., 2016).

2 Salzman, J., Gawad, C., Wang, P. L., Lacayo, N. & Brown, P. O. Circular RNAs are the predominant transcript isoform from hundreds of human genes in diverse cell types. *PLoS One* **7**, e30733, doi:10.1371/journal.pone.0030733.

3 Jeck, W. R. *et al.* Circular RNAs are abundant, conserved, and associated with ALU repeats. *RNA* **19**, 141-157, doi:10.1261/rna.035667.112.

4 Zhang, X. O. *et al.* Complementary sequence-mediated exon circularization. *Cell* **159**, 134-147, doi:10.1016/j.cell.2014.09.001 (2014).

5 Chen, C. Y. & Sarnow, P. Initiation of protein synthesis by the eukaryotic translational apparatus on circular RNAs. *Science* **268**, 415-417 (1995).

6 Wang, Y. & Wang, Z. Efficient backsplicing produces translatable circular mRNAs. *RNA* **21**, 172-179, doi:10.1261/rna.048272.114.

7 Legnini, I. *et al.* Circ-ZNF609 Is a circular RNA that can be translated and functions in myogenesis. *Mol Cell* **66**, 22-37 e29, doi:10.1016/j.molcel.2017.02.017 (2017).

8 Pamudurti, N. R. *et al.* Translation of circRNAs. *Mol Cell* **66**, 9-21 e27, doi:10.1016/j.molcel.2017.02.021 (2017).

9 Yang, Y. *et al.* Extensive translation of circular RNAs driven by N(6)-methyladenosine. *Cell Res* **27**, 626-641, doi:10.1038/cr.2017.31 (2017).

10 Li, X. *et al.* Coordinated circRNA biogenesis and function with NF90/NF110 in viral infection. *Mol Cell* **67**, 214-227 e217, doi:10.1016/j.molcel.2017.05.023 (2017).

11 Fairbrother, W. G., Yeh, R. F., Sharp, P. A. & Burge, C. B. Predictive identification of exonic splicing enhancers in human genes. *Science* **297**, 1007-1013, doi:10.1126/science.1073774 (2002).

12 Wang, Y., Ma, M., Xiao, X. & Wang, Z. Intronic splicing enhancers, cognate splicing factors and context-dependent regulation rules. *Nat Struct Mol Biol* **19**, 1044-1052, doi:10.1038/nsmb.2377 (2012).

13 Weingarten-Gabbay, S. *et al.* Systematic discovery of cap-independent translation sequences in human and viral genomes. *Science* **351**, doi:10.1126/science.aad4939 (2016).

14 Gritsenko, A. A. *et al.* Sequence features of viral and human Internal Ribosome Entry Sites predictive of their activity. *PLoS Comput Biol* **13**, e1005734, doi:10.1371/journal.pcbi.1005734 (2017).

15 Mitchell, S. A., Spriggs, K. A., Coldwell, M. J., Jackson, R. J. & Willis, A. E. The Apaf-1 internal ribosome entry segment attains the correct structural conformation for function via interactions with PTB and unr. *Mol Cell* **11**, 757-771, doi:10.1016/s1097-2765(03)00093-5 (2003).

16 Paz, I., Kosti, I., Ares, M., Jr., Cline, M. & Mandel-Gutfreund, Y. RBPmap: a web server for mapping binding sites of RNA-binding proteins. *Nucleic Acids Res* **42**, W361-367, doi:10.1093/nar/gku406 (2014).

17 Dominguez, D. *et al.* Sequence, structure, and context preferences of human RNA binding proteins. *Mol Cell* **70**, 854-867 e859, doi:10.1016/j.molcel.2018.05.001 (2018).

18 Van Nostrand, E. L. *et al.* A large-scale binding and functional map of human RNA-binding

proteins. *Nature* **583**, 711-719, doi:10.1038/s41586-020-2077-3 (2020).

19 Dresios, J., Chappell, S. A., Zhou, W. & Mauro, V. P. An mRNA-rRNA base-pairing mechanism for translation initiation in eukaryotes. *Nat Struct Mol Biol* **13**, 30-34, doi:10.1038/nsmb1031 (2006).

20 Chappell, S. A., Edelman, G. M. & Mauro, V. P. A 9-nt segment of a cellular mRNA can function as an internal ribosome entry site (IRES) and when present in linked multiple copies greatly enhances IRES activity. *Proc Natl Acad Sci U S A* **97**, 1536-1541, doi:10.1073/pnas.97.4.1536 (2000).

21 Owens, G. C., Chappell, S. A., Mauro, V. P. & Edelman, G. M. Identification of two short internal ribosome entry sites selected from libraries of random oligonucleotides. *Proc Natl Acad Sci U S A* **98**, 1471-1476, doi:10.1073/pnas.98.4.1471 (2001).

22 Zeenko, V. & Gallie, D. R. Cap-independent translation of tobacco etch virus is conferred by an RNA pseudoknot in the 5'-leader. *J Biol Chem* **280**, 26813-26824, doi:10.1074/jbc.M503576200 (2005).

23 Nicholson, R., Pelletier, J., Le, S. Y. & Sonenberg, N. Structural and functional analysis of the ribosome landing pad of poliovirus type 2: in vivo translation studies. *J Virol* **65**, 5886-5894, doi:10.1128/JVI.65.11.5886-5894.1991 (1991).

24 Chappell, S. A. & Mauro, V. P. The internal ribosome entry site (IRES) contained within the RNA-binding motif protein 3 (Rbm3) mRNA is composed of functionally distinct elements. *J Biol Chem* **278**, 33793-33800, doi:10.1074/jbc.M303495200 (2003).

25 Panek, J., Kolar, M., Vohradsky, J. & Shivaya Valasek, L. An evolutionary conserved pattern of 18S rRNA sequence complementarity to mRNA 5' UTRs and its implications for eukaryotic gene translation regulation. *Nucleic Acids Res* **41**, 7625-7634, doi:10.1093/nar/gkt548 (2013).

26 Pisarev, A. V., Kolupaeva, V. G., Yusupov, M. M., Hellen, C. U. & Pestova, T. V. Ribosomal position and contacts of mRNA in eukaryotic translation initiation complexes. *EMBO J* **27**, 1609-1621, doi:10.1038/emboj.2008.90 (2008).

27 Matsuda, D. & Mauro, V. P. Base pairing between hepatitis C virus RNA and 18S rRNA is required for IRES-dependent translation initiation in vivo. *Proc Natl Acad Sci U S A* **111**, 15385-15389, doi:10.1073/pnas.1413472111 (2014).

28 Malygin, A. A., Kossinova, O. A., Shatsky, I. N. & Karpova, G. G. HCV IRES interacts with the 18S rRNA to activate the 40S ribosome for subsequent steps of translation initiation. *Nucleic Acids Res* **41**, 8706-8714, doi:10.1093/nar/gkt632 (2013).

29 Meng, Z., Jackson, N. L., Shcherbakov, O. D., Choi, H. & Blume, S. W. The human IGF1R IRES likely operates through a Shine-Dalgarno-like interaction with the G961 loop (E-site) of the 18S rRNA and is kinetically modulated by a naturally polymorphic polyU loop. *J Cell Biochem* **110**, 531-544, doi:10.1002/jcb.22569 (2010).

#### Experimental procedures

##### Plasmid library construction and screening

In order to screen for cap-independent translation initiation elements in the human transcriptome, the previously described pcircGFP reporter [1] was modified to generate stable cell lines with the Flp-in system. The DNA fragment that contains the exon of split GFP and complementary introns were inserted into pcDNA5/FRT/TO plasmid. To increase ligation efficiencies and to reduce the frequencies of restriction sites, the AscI and NotI restriction sites were inserted between the start codon and stop codon, generating pcircGFP-IREScreen.

To produce the library of human transcriptome fragments, we extracted total RNA from HeLa cells with TRIzol reagent (Invitrogen) according to the manufacturer′s instructions. Total RNAs (5 μg) were treated with DNase I (Promega RQ1 RNase-free DNase) to remove genomic DNA, and then fragmented by heating at 85°C for 6 min in fragmentation buffer (100 mM Tris, pH 8.0, 2 mM $MgCl_2$). The RNA fragments were reverse-transcribed to generate strand-specific double-strand cDNAs (ds-cDNAs), and the ds-cDNAs were then end-repaired, A-tailed and ligated to illumina adaptors (see details in Extended data Table 4) using a KAPA stranded mRNA-seq kit (KAPA biosystems) according to the manufacturer's instructions. To reduce abundant ds-cDNAs derived from rRNA and housekeeping genes, the ds-cDNAs were normalized by duplex-specific nuclease (DSN from evrogen) treatment. 500 ng of ds-cDNAs in hybridization buffer (50 mM HEPES, pH7.5, 500mM NaCl) were denatured at 98°C for 2 min and annealed by incubating at 68°C for 5 h. Then, the cDNA sample was treated with DSN at 68°C for 30 min. The DSN-treated cDNA sample was purified using SPRI beads and amplified with KAPA HiFi DNA polymerase (KAPA biosystems). To generate the plasmid library, the amplified DSN-treated cDNA sample was digested with AscI and NotI (NEB) and ligated into AscI and NotI digested pcircGFP-IREScreen plasmid. The ligation product was transformed into ElectroMax DH-5α (Invitrogen), and 4,500,000 *E. coli* clones were obtained. The resulting library (denoted the IRESceen plasmid library) was extracted using a QIAGEN Plasmid Mega Kit.

To avoid false discoveries due to transcription readthrough of the plasmid and increase the accuracy of the screen, we used the Flp-In system to generate a stable cell line library that contains one copy of each inserted fragment in each cell. The IRESceen plasmid library and pOG44 (ratio 1:9) were transfected into Flp-In™-293 cells (Invitrogen) (50 μg per 15 cm dish) using lipofectamine 2000 (Invitrogen). To obtain enough stable cell clones, four batches of transfection (each with 20 dishes) were conducted. Each dish was split into four at 24 h after transfection, and the cells were selected by hygromycin (100 μg /mL) for 2 weeks.

To identify the inserted fragments with IRES activity, we used FACS to sort the green cells. Two days before performing FACS, the hygromycin was depleted from the culture medium. To select the singlets, SSC-A *vs.* FSC-A was used to select Flp-In™-293 cells (excluding very small and very large particles). Two round selections of singlets were used by measuring SSC-W *vs.* FSC-H and FSC-W *vs.* FSC-H. Then FITC-A *vs.* PE-A was used to select GFP-positive cells. RNAs were subsequently extracted, and sequencing libraries were generated using RT-PCR. RNA-seq was performed with Hiseq 2500.

**Cell cultures and Transfection**

HEK293 and SH-SY5Y cell lines were cultured in DMEM (high glucose, Hyclone) medium containing 10% fetal bovine serum (FBS, Hyclone). The HCT116 cell line was cultured in McCoy's 5A (Gibco) medium containing 10% fetal bovine serum (FBS, Hyclone). To transiently transfect plasmids into cells, 2 μg of mini-gene reporters were transfected into cells in a 6-well plate, using lipofectamine 3000 (Invitrogen) according to the manufacturer's instructions. After 48 h, cells were collected for further analysis of RNA and protein levels.

**Semi-quantitative RT-PCR and real-time PCR**

Total RNAs were isolated from transfected cells with TRIzol reagent (Invitrogen) according to the manufacturer's instructions. Total RNAs (1 μg) were treated by gDNA eraser to remove genomic DNA, then reverse-transcribed with a PrimeScript RT Reagent kit with gDNA Eraser (TaKaRa) using a mixture of oligo(dT) and random hexamer primers (following the manufacturer's instructions). Then, RT products (1 μl) were used as the template for PCR amplification (25 cycles of amplification). PCR products were separated on 10% polyacrylamide gel electrophoresis (PAGE) gels, visualized by staining with SYBR Green I (Thermo Scientific) and scanned using ChemiDoc Touch Image system (BioRad).

**Western blot**

Cells were lysed in RIPA buffer containing a protease inhibitor cocktail (Roche), and the total cell lysates were resolved on a 4-20% ExpressPlus™PAGE Gel (GeneScript). The following antibodies were used: GFP antibody (Clontech: 632381), Flag epitope antibody (Sigma: F1804-1MG), HA epitope antibody (CST: 3724), DHX29 antibody (Bethyl laboratories: A300-751A), HIF1A antibody (CST: 14179), and Renilla Luciferase antibody (abcam: ab187338). Antibodies were diluted by 1:2000. HRP-conjugated GAPDH antibody (ABclonal: AC035), and HRP-conjugated ACTB antibody (ABclonal: AC028) were diluted by 1:10000. The HRP-linked secondary antibodies (CST: anti-mouse IgG 7076S, anti-rabbit IgG 7074S) were used at a 1:2000 dilution and the blots were visualized with the ECL reagents (Bio-Rad).

**Dual Luciferase assay**

The dual luciferase reporter (psi-CHECK2) was obtained from Promega. The optional *cis*-elements were inserted into the 5′UTR (such as hairpin structures, uORFs or 5′UTRs of endogenous genes) or into the 3′UTR (such as various CiTIs, 3′UTR of

HIF1A, or MS2 sites) of the Rluc gene to test their effects on mRNA translation, and the Fluc gene was used as an internal control. These modified reporters were transfected into HEK293, SH-SY5Y or HCT116 cells in a 24-well plate (100 ng reporter per well). At 36 h after transfection, cells were collected and lysed in the passive lysis buffer (Promega). Protein expression was measured via luminescence that was determined by Bio-Tek synergy H1 using the dual-luciferase reporter assay system (Promega), and RNA expression was determined by RT-qPCR (Roche LC480).

**Gene knockdown with lentiviral shRNA**

The pairs of oligos denoted DHX29-sh1 (the siRNA region sequence: CGTGTACCTTTGGAGGAATTA), DHX29-sh2 (the siRNA region sequence: CCTAAGTATCAGAAACTTCTA), or HIF1A-sh (the siRNA region sequence: TGCTCTTTGTGGTTGGATCTA) were 5′-phosphorylated using T4 Polynucleotide Kinase (NEB) and then annealed pair-wisely. The annealed oligos were subsequently ligated into the AgeI/EcoRI digested pLKO.1 vector. Then, the resulting pLKO.1-scramble plasmid or pLKO.1-shRNA plasmids were transfected into HEK293 cells with psPAX2 and pMD2.G at a ratio of 4:3:1 using lipofectamine 3000 (Thermo Fisher). The resulting virus was collected at 48h after transfection. The SH-SY5Y cells were infected by the control or shRNA lentivirus for 48h, followed by 5 μg/ml puromycin selection.

**RNP affinity purification and mass spectrometry**

Uncapped RNAs containing the CiTI or control sequences that were fused with a glmS ribozyme followed by three MS2 binding sites were *in vitro* transcribed from linearized plasmids using T7 RNA polymerase (homemade), in the presence or absence of $^{32}$P-UTP. To facilitate the detection of the RNA and provide sufficient amounts for affinity purification, radiolabeled and non-radiolabeled RNAs were mixed together. The resulting RNAs (2 nmol) were incubated with MS2–MBP protein (6 nmol) in 150 μl of binding buffer (16 mM HEPES (pH 7.6), 4.5 mM $Mg(OAc)_2$, 125 mM KOAc, 8 μg/ml tRNA, 0.8 mM ATP and 0.1 mM GTP) for 1 hr on ice. Four ml of HeLa cytoplasmic extract were mixed with 6 ml of binding buffer and pre-incubated for 5 min at room temperature. The pre-incubated cytoplasmic extract was subsequently added to the RNA and MS2–MBP mixture, and incubated at room temperature for 10 min. The mixtures were loaded onto amylose columns (500 μl bed volume for each RNA) pre-equilibrated with 2.5 ml binding buffer at 4°C and washed 3X with binding buffer. To activate self-cleavage of the glmS ribozyme, 250 μl elution buffer (binding buffer containing 1 mM GlcN6P) was added to each column and incubated for 20 min at room temperature. The RNP complexes were eluted dropwise with 1.2 ml elution buffer and collected in 8 fractions. The amount of RNP complexes in each fraction was determined by quantitating the amount of $^{32}$P -labeled RNA present by Cherenkov counting. Proteins in the third eluate (containing the peak of $^{32}$P -labeled RNA) were analyzed by SDS-PAGE. Proteins were in-gel digested with trypsin overnight and the resulting peptides were separated on a C18 column using an UltiMate3000 (Dionex) ultrahigh performance liquid chromatography system, and analyzed by electrospray ionization mass spectrometry using a Thermo Scientific Q Exactive HF mass spectrometer. Proteins bound to the CiTI or control sequences were identified by searching fragment spectra against the Uniprot database using Mascot as a search engine. Proteins copurifying with all of the CiTI-containing RNAs, and additionally exhibiting a >2-fold enrichment in peptide counts (relative to the control RNA), were analyzed further.

***In vivo* icSHAPE**

The icSHAPE experiments were performed in HEK293 cells that were transfected with the uAUG-hp35 reporter or uAUG-hp35-CiTI reporter as previously described with some modifications [2-4]. Briefly, cells were dissociated with trypsin and washed with PBS. The harvested cells were then incubated with 0.1M NAI-N3 (used to label flexible RNA bases) or DMSO (as a control) at 37°C for 5 min with rotation. The labeled and control cells were centrifuged at 2500g for 1 min at 4°C, and washed by PBS. Then total RNAs were extracted from cells with TRIzol reagent (Invitrogen) according to the manufacturer's instructions, and were treated by DNase I to remove genomic DNA. The DNase I treated RNAs were purified using an RNA Clean & Concentrator kit (ZYMO Research), and mRNAs were enriched using an NEBNext Poly(A) mRNA Magnetic Isolation Module (NEB).

The *in vivo* modified mRNA and control unmodified mRNA samples were combined with 2 μL 1.85mM DIBO-biotin (ThermoFisher) solution and 1 μL

RNasin® Ribonuclease Inhibitor (Promega) and incubated on a Thermomixer at 37°C for 2 h. The resulting RNAs were fragmented in buffered zinc solution (Ambion RNA fragmentation reagents). To repair the ends of the fragmented RNAs, samples were incubated in 10 μL end repairing mix (70 mM Tris 7.0, 18 mM $MgCl_2$, 5 mM DTT, 4 U/μL RNasin® Ribonuclease Inhibitor, 0.1 U/μL FastAP (Life Technology), 2 U/μL T4 PNK(NEB)) at 37°C for 1 h. Then 10 μL ligation mix (5 mM DTT, 1.25 μM 5′ adenylylated and 3′-blocked linker (3′-biotin for unmodified, 3′-NH3 for modified), 0.66 U/μL T4 RNA ligase (NEB, M0437M), 15% PEG8000, 1X RNA ligase buffer) were added to each sample and incubated at 25°C for 3 h. The resulting samples were purified using an RNA Clean & Concentrator kit (ZYMO Research), and the purified RNAs were incubated at 30°C for 90 min in 10 μL reaction buffer containing 1μL FastAP, 0.2μL SSB (Promega), 0.8 μL RNasin® Ribonuclease Inhibitor and 1 μL 5′ Deadenylase (NEB) in 1X NEB buffer 2 (NEB). To remove 3′ adaptor contamination, 1 μL RecJf (NEB) was added and the reaction was incubated at 37°C for an additional 1 h. The subsequent procedures were the same as described for the standard protocol [4].

The icSHAPE-pipe [5] was used to align the deep sequencing data to the custom reference genome, which includes the GRCh37 human reference genome with gencode v32lift37 annotation and an additional reporter sequence (uAUG-hp35). Subsequently, the icSHAPE signal at each position was calculated as the $Modified_{signal}$ / $Untreated_{signal}$ from bedGraph files. RNA secondary structures were reconstructed by RNAfold based on their icSHAPE signals.

**Polysome profiling and sequencing**

Cultured cells were pre-treated with 100 μg/mL cycloheximide for 5 min at 37 °C and washed with ice-cold PBS containing 100 μg/mL cycloheximide. Cells were then lysed in polysome lysis buffer (400 mM KCl, 10 mM HEPES (pH 7.4), 5 mM $MgCl_2$, 1 mM DTT, 100 μg/mL cycloheximide, 0.5% Triton$^{TM}$ X-100, 0.5% deoxycholate, 1X protease inhibitor cocktail and 50 U/ml RNasin) by incubating for 10 min on ice. Cell debris was removed by centrifugation at 14,000 rpm in an T15A61 rotor (HITACHI CT15E) for 10 min at 4°C, and the supernatant was loaded onto 10 mL continuous 10-50% sucrose gradients containing 400 mM KCl, 10 mM HEPES (pH 7.4), 5 mM $MgCl_2$, 100 μg/mL cycloheximide, 1X protease inhibitor cocktail and 50 U/ml RNasin. The samples were centrifuged at 4 °C for 3 h at 35,000 × rpm in an SW41 rotor (Beckman). Gradient fractions were collected using a Brandel Fractionation System, and the polysome profiles of the gradients were generated using an Isco UA-6 ultraviolet detector.

An *in vitro* synthesized Fluc RNA (1 ng per 500 mL fraction) was added to each polysome fraction as a spike-in reference to quantify the relative RNA expression level in each polysome fraction. To identify the genes with differential translation levels through high-throughput sequencing, the free fractions (free, 40s and 60s fractions), light polysome fractions (fractions with 2-4 ribosomes) or heavy polysome fractions (fractions with more than 4 ribosomes) were pooled together in an equal volume, respectively. Total RNAs were then extracted from these fractions and used for the RNA-seq library preparation using a KAPA stranded RNA-seq kit with RiboErase. To quantify the HIF1A or ACTB mRNA level by RT-qPCR, the total RNA was extracted from each fraction with TRIzol reagent, and the resulting RNA was reverse transcribed for the quantification of gene expression using qPCR (Roche LC480).

**Deletion of 3′-CITI in HIF1A using CRISPR-cas9**

To delete the 3′-CiTI of HIF1A gene, two guide RNAs were designed: gRNA1 (ATGATGCTACTGCAATGCAA), and gRNA2 (TCCAGGTTTAACAATTTCAT). The oligos of the gRNAs were 5′-phosphorylated by T4 Polynucleotide Kinase (NEB) and then annealed together. The annealed oligos were further ligated into the BbsI-digested eSpCAS9 vector. The resulting plasmids (eSpCAS9-gRNA1 and eSpCAS9-gRNA2) were transfected into SH-SY5Y cells at 1:1 ratio using Lipofectamine 3000. At 2 days after transfection, a part of the pooled cell population was collected to determine the efficiency of CRISPR genome digestion. Then, the single cells were seeded into 96 wells, and the genotype of each cell population was determined by PCR at 10 days after seeding. The positive cell clones were used for further analysis.

**Cell proliferation assay**

One day before the cell proliferation experiment, cells were seeded into 96 well plates (1,000 cells each well). 12 h later, the relative starting cell numbers were determined using a Cell Counting Kit-8 (Dojindo Molecular Technologies). Subsequently, the relative cell numbers were measured every 24 hours. For the hypoxia treatment condition, the cells were cultured under hypoxia conditions (1% $O_2$) at 12 h after the cell seeding.

**Xenograft tumor model**

The mouse experiments were approved by the Institutional Animal Care and Use Committee at the Shanghai Research Centre for Model Organisms (IACUC NO: 2018-0004). The SH-SY5Y (WT, Del-C1, or Del-C2) cells (1.0 x $10^7$ cells in 100 μl PBS per mouse) were injected into nude mice (nine mice for each cell clone). Starting one week after injection, the tumor size was measured twice a week. Mice were then euthanized 4 weeks after tumor cell inoculation, and the transplanted tumors were collected for tumor weight measurement.

**Computational identification and analyses of CiTI sites**

High-throughput sequencing reads (starting and pre-sorting libraries: 51bp paired-end reads; post-sorting library: 101bp paired-end reads) were trimmed using Cutadapt (version 1.16) to remove Illumina adaptor sequences and subsequently aligned to human GRCh37 genome with STAR (version 2.4.2a) using the additional parameters: '--outFilterMultimapNmax 1 --outFilterScoreMin 10 --outFilterMismatchNmax 3'. The resulting paired-end reads were used to define the regions of RNA fragments. A large fraction (84%) of the paired-end reads in the post-sorting library overlapped with each other. To further analyze the features of these RNA fragments within protein coding genes, we reconstructed the full-length sequences using the paired-end reads. The sequences of overlapping reads were directly regarded as the full-length CiTI fragments. The gaps between non-overlapping, paired-end reads were filled using the sequences of annotated transcripts with the longest exon in these regions (GENCODE v27lift37), and the resulting sequences were regarded as the full-length RNA fragments. The RNA fragments in the post-sorting library were used to define the sites (termed as CiTIs) which contained at least one fragment.

To estimate the rRNA ratio in starting, pre-sorting, or post-sorting libraries, RNA-seq reads were uniquely aligned to rDNA sequences by Bowtie (bowtie version 1.1.2). Gene expression levels were quantified using RSEM (v1.3.1, --gzipped-read-file --paired-end --strand-specific --calc-ci --calc-pme --time --keep-intermediate-files --star). To assess the conservation levels of different regions of mRNA, PhastCons46 scores computed from genome alignments of 46 vertebrates including humans (hg19) were extracted from the PhastCons46 track in the UCSC Genome Browser. Metagene plots were constructed by normalized read density around the start and stop codons of each gene (i.e, read depth at each position divided by the number of the covered transcripts).

To compare the CiTIs with the known IRESs, we used two datasets of experimentally studied IRESs: a known IRES collection in IRESite database (http://www.iresite.org/) (Mokrejs et al., 2010) and the identified IRESs in previous screen with the bicistronic reporter (Weingarten-Gabbay et al., 2016) (the raw data was provided by Weingarten-Gabbay Shira, and the thresholds were set as suggested: fluorescence score > 600, promoter activity < 0.2, and splicing activity > -2.5).

**Analysis of sequences complementary to 18S rRNA**

Complementarity to the 18S rRNA was defined by base-pairing between the candidate RNAs and 18S rRNA with an at least 7nt consecutive match. To calculate the enriched regions in 18S rRNA that paired with different types of CiTIs, the number of paired sequences (with CiTIs) at each position of 18S rRNA was divided by the total number of CiTIs. As background controls, we also calculated the complementarity of different mRNA regions (5′UTR, CDS, and 3′UTR) to the 18S rRNAs

**Motif analysis**

The overlapping hexamers were extracted from CiTI or control sequences, and the occurrences of each hexamer were determined for the CiTI or control sequences. The enrichment score of each 6-mer between the CiTI and control was calculated using Z-test. Hexamers with a score larger than 4 were defined as enriched motifs. The resulting elements were aligned with CLUSTALW2 [6] to generate consensus motifs, which were plotted by Weblogo3.

**Identification of dORF-coded proteins**

Using an open search engine, pFind (v3.1.3) [7], we searched the published human comprehensive proteome datasets [8,9] against a customized database containing all UniProt human proteins and the potential dORF-coded proteins previously identified by Ribo-seq [10,11]. We selected positive mass spectra using the following thresholds: q ≤0.01, missed cleavage sites ≤3, allowing only common modifications (cysteine carbamidomethylation, oxidation of methionine, protein N-terminal acetylation, pyro-glutamate formation from glutamine, and phosphorylation of serine, threonine, and tyrosine residues).

**Analysis of protein-protein interaction network**

To perform the protein interaction analysis, we selected the genes containing highly confident CiTIs (read depth ≥3), and searched STRING with the parameters of medium confidence, considering all interaction evidence and discarding disconnected nodes. The resulting networks were clustered using the MCODE tool in the Cytoscape package. The function of each cluster was annotated using the DAVID Gene ontology tool.

**Data availability**

The raw sequencing data and filtered datasets are publicly available from NODE (assess number: OEP001285) and NCBI (assess number: PRJNA680213). The customized scripts for data analyses are also deposited in GitHub: https://github.com/rnasys/CiTI.

### Extended data figures and figure legends

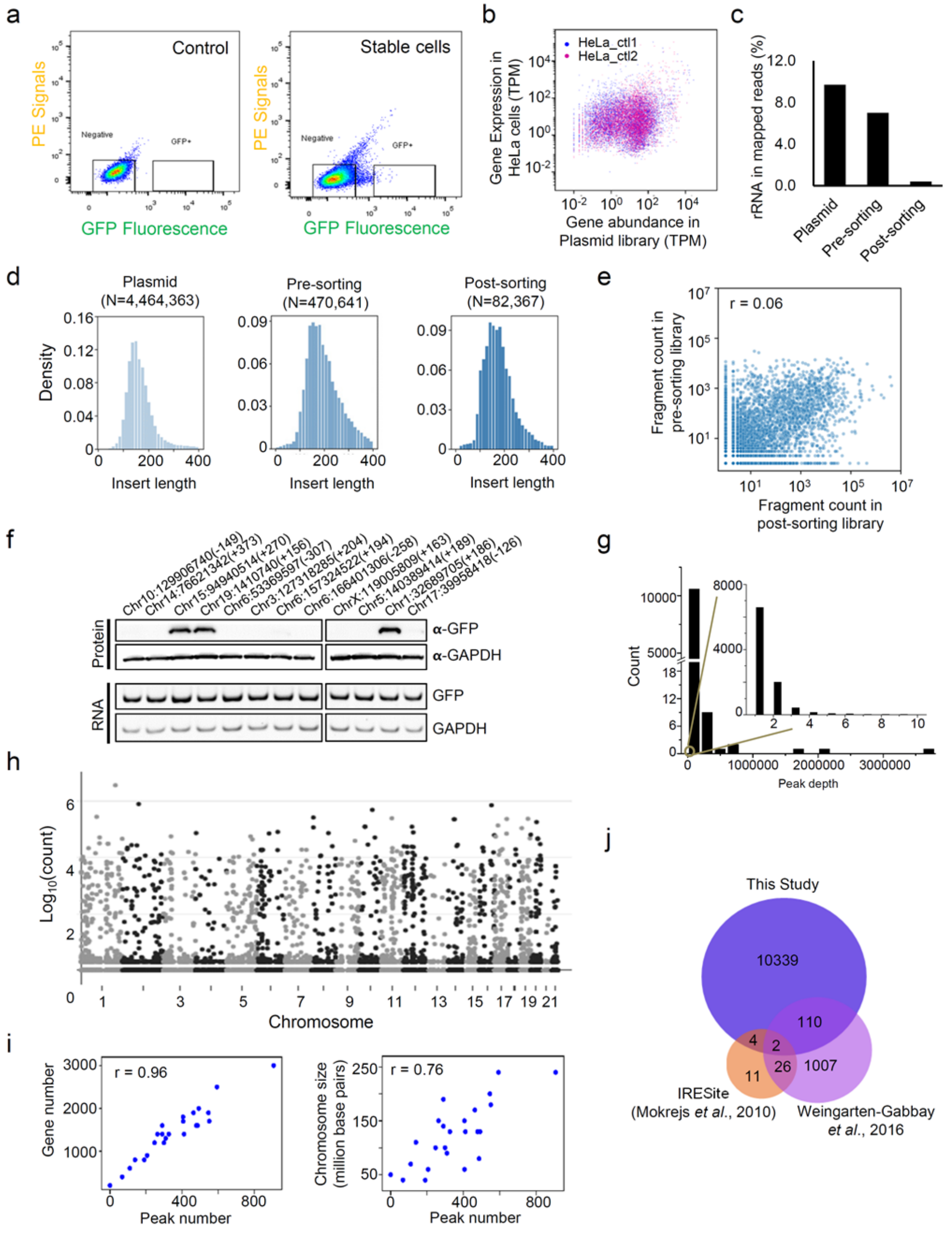


**Extended data figure 1. The screen for human, endogenous *cis*-elements involved in cap-independent translation. a**, The gate setting of FACS by which the cells with high GFP fluorescence were sorted. **b**, The correlation between the gene expression of

HeLa cells [12] and the fragment abundance in the plasmid library. **c**, Ribosomal RNA abundance in plasmid, pre-sorting and post-sorting libraries. **d**, Length distribution of the inserted sequences within the plasmid, pre-sorting and post-sorting libraries. **e**, The abundance of inserted sequences within the circRNAs in pre-sorting and post-sorting cells (Spearman correlation coefficient, r = 0.06). **f**, Validation of the random selected fragments (indicated by their starting position in the human genome and offset, +: plus-strand, -: minus-strand, see sequences in Extended data table 4) for cap-independent translation activation using circRNA. CircRNA reporters with different inserted fragments were transfected into HEK293 cells, and the translation products were detected by western blot. **g**, Read depth distribution of newly identified CiTIs. **h**, Distribution of the CiTIs across human chromosomes. **i**, Correlation between the abundance of CiTIs and gene numbers of each chromosome, or the size of each chromosome. **j**, Overlap of the cap-independent sequences derived from our screen and previous studies [13,14].

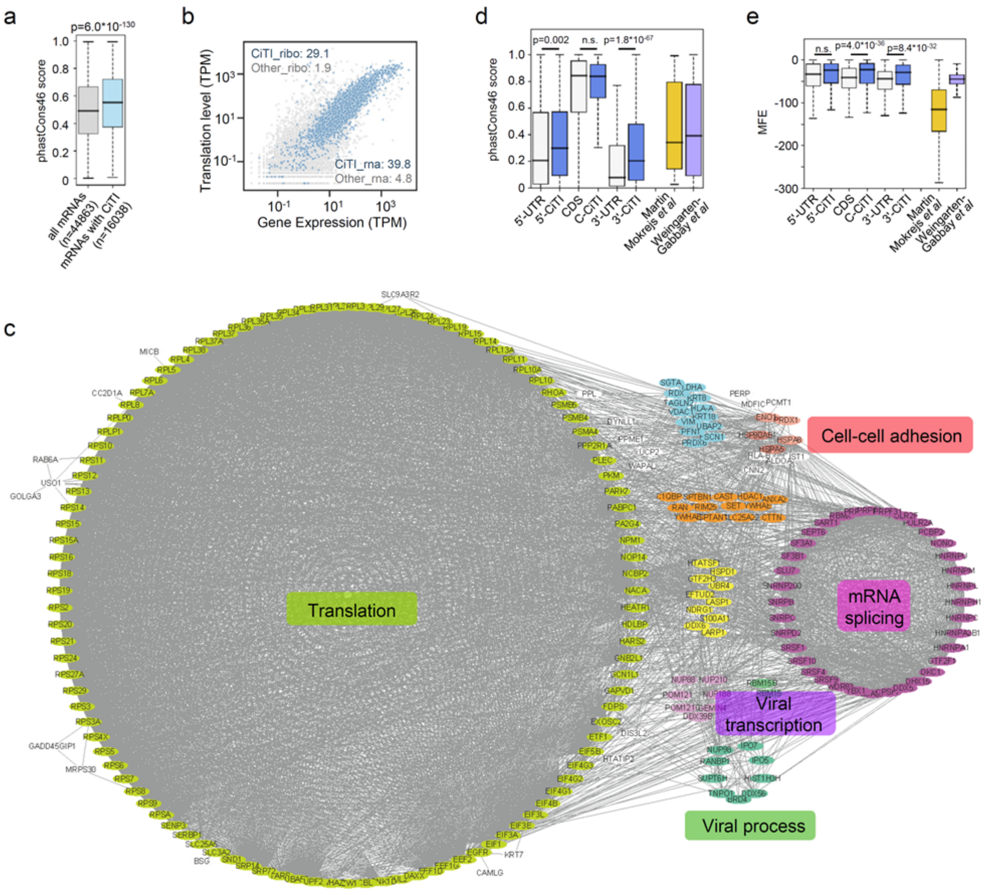


**Extended data figure 2. Distinct features of human endogenous CiTIs. a**, Sequence conservation of the CiTI-containing genes (p value was calculated by unpaired two-sided Mann–Whitney U test). **b**, RNA expression or translation efficiency of mRNAs in cultured cells. RNA expression was determined by RNA-seq, and the translation efficiency was measured using Ribo-seq [15]. Blue dots represent the CiTI-containing genes. The median expression levels and translation levels were indicated as CiTI-rna, Other-rna, CiTI-ribo and Other-ribo, respectively. **c**, Protein-protein interactions of the genes containing highly confident CiTIs (read depth ≥3). The disconnected nodes were hidden. **d**, Sequence conservation and **e**, Predicted minimal free energy (MFE) of the CiTIs within different gene regions and the cognate control regions. The known IRESs from the IRESite collection [13,14] or a previous screen [13,14] were also included as controls. P values were calculated using the unpaired two-sided Mann-Whitney U test.

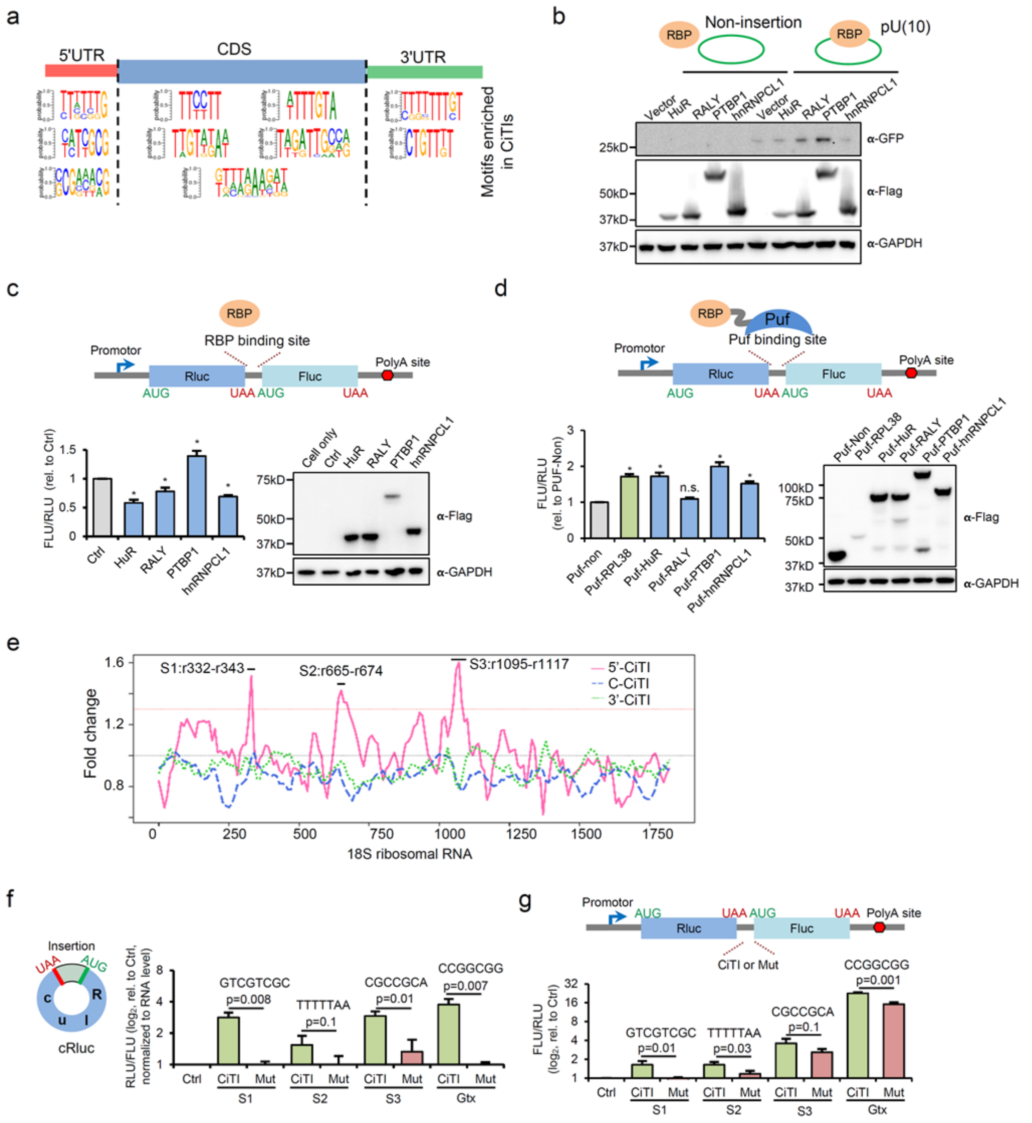


**Extended data figure 3. Short motifs of human endogenous CiTIs. a**, Enriched motifs in CiTIs located in the 5′UTR, CDS, or 3′UTR regions. **b-d**, *Trans*-acting factors promote cap-independent translation. **b**, The circRNA or **c**, bicistronic reporters with a poly-U motif were co-transfected together with the plasmids coding for *trans*-acting factors into HEK293 cells. The translation products (GFP) of circRNAs were judged by Western blot, and the translation products (luciferases) of bicistronic reporters were measured by luminescence assay (n = 3, mean ± SD, * indicated p<0.05 by Student's t-test). **d**, Cap-independent translation is promoted through binding of *trans*-acting factors. Bicistronic reporters containing Puf binding sites were co-transfected with the reporters encoding the fusion proteins with RBPs and a Puf domain into HEK293 cells, and the translation products were examined using luminescence assays (n = 3, mean ± SD, * indicated $p<0.05$, n.s. indicated no significant difference by Student's t-test). **e**, Complementarity of different 18S rRNA regions to the CiTIs, which was plotted as relative fold change to the cognate mRNA background regions. Three distinct regions of 18S rRNAs (S1, S2 and S3) that significantly paired with 5′-CiTIs are indicated by horizontal lines. **f-g**, Cap-independent translation activity of 5′-CiTIs with 18S rRNA complementary motifs. The 5′-CiTIs with 18S rRNA complementary motifs (or the mutated CiTIs as controls) were inserted into the circRNA (**f**) or bisctronic reporters (**g**), which were transfected into HEK293 cells to measure the translation products by luminescence assay 36 hours after transfection (n = 3, mean ± SD, Student′s t-test).

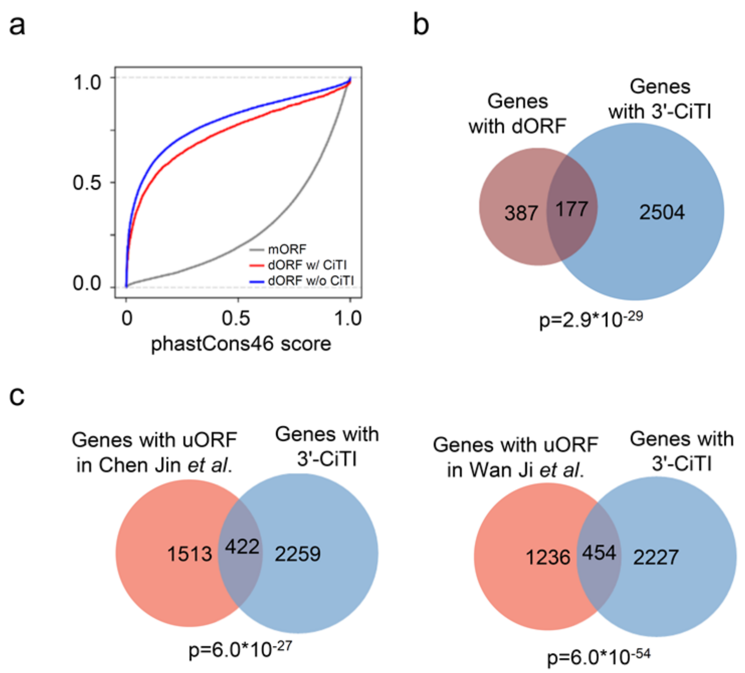


**Extended data figure 4. Analysis of downstream ORFs in the 3′UTR and features in the 5′UTR of 3′-CiTI containing genes. a**, Accumulation curve for the sequence conservation score of main ORFs (mORF), and predicted dORFs with or without 3′-CiTIs. The dORFs in 3′UTRs were predicted with the ORFs longer than 60 nt. **b**, Overlap of the genes with 3′-CiTIs and dORFs. The dORFs were obtained from published results [10,11] (p value was calculated by hypergeometric test). **c**, Overlap of the genes with uORFs and 3′-CiTIs. The uORFs were obtained from published results [16,17] (p value was calculated by hypergeometric test).

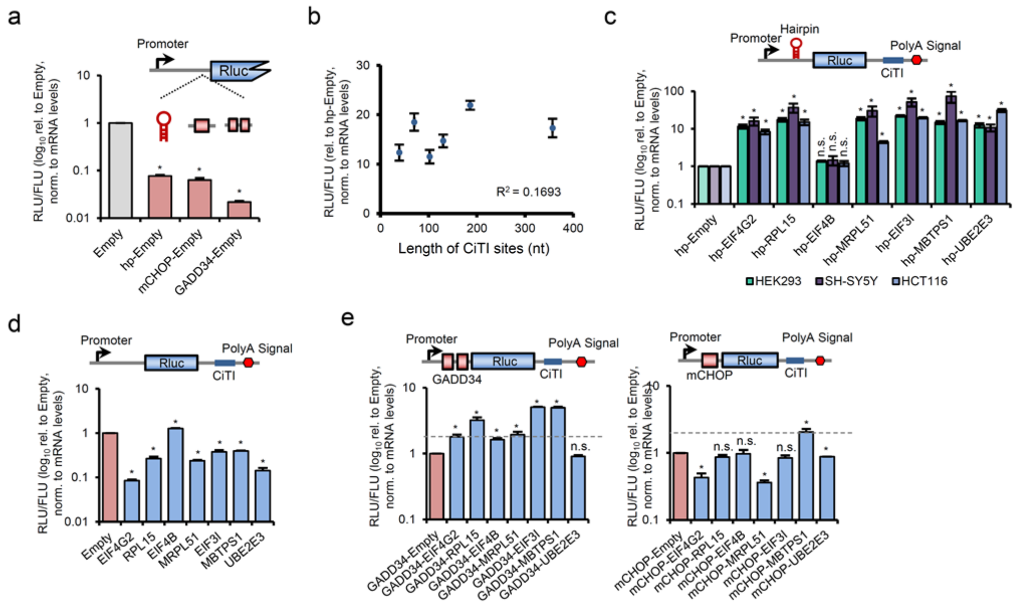


**Extended data figure 5. The diverse activities of 3′-CiTIs in different reporter systems. a**, The dual-luciferase reporters containing different 5′UTRs with strong structures or uORFs were transfected into HEK293 cells, and the translation efficiency of Rluc was determined by the relative abundance of proteins (*via* luminescence assay) *vs*. RNAs (*via* qPCR) normalized to the control Fluc (n = 3, mean ± SD, * indicated p<0.05 by Student's t-test). **b**, No correlation between the length of CiTIs and their activities in reporters with structured 5′UTRs. **c**, The dual-luciferase reporters containing structured 5′UTRs and various CiTIs in their 3′UTR were transfected into SH-SY5Y and HCT116 cells. The same experiments and analyses were performed as described in **panel a** (n = 3, mean ± SD, * indicated p<0.05, n.s. indicated no significant difference by Student's t-test). **d**, The various CiTIs were inserted into 3′UTR regions of the Rluc gene without inserted regulatory elements in the 5′UTRs. The same analyses were performed as described in **panel a** (n = 3, mean ± SD, * indicated p<0.05 by Student's t-test). **e**, Different CiTIs were inserted into 3′UTR regions of the Rluc gene containing uORFs in their 5′UTRs. The same analyses were performed as described in **panel a** (n = 3, mean ± SD, * indicated p<0.05, n.s. indicated no significant difference by Student's t-test).

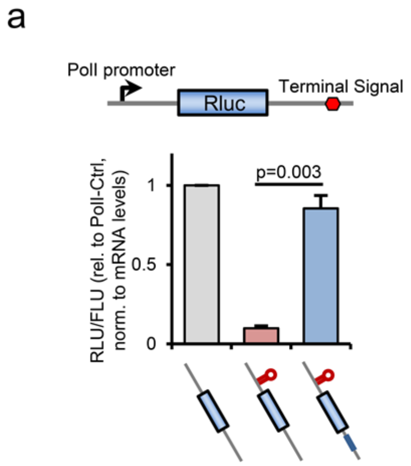


**Extended data figure 6. Validation of CiTI activity using PolI reporter. a**, Examination of 3′-CiTI activity using reporters with a PolI promoter and terminator, which will generate mRNAs without a 5′-cap and poly-A tail (n = 3, mean ± SD, Student′s t-test).

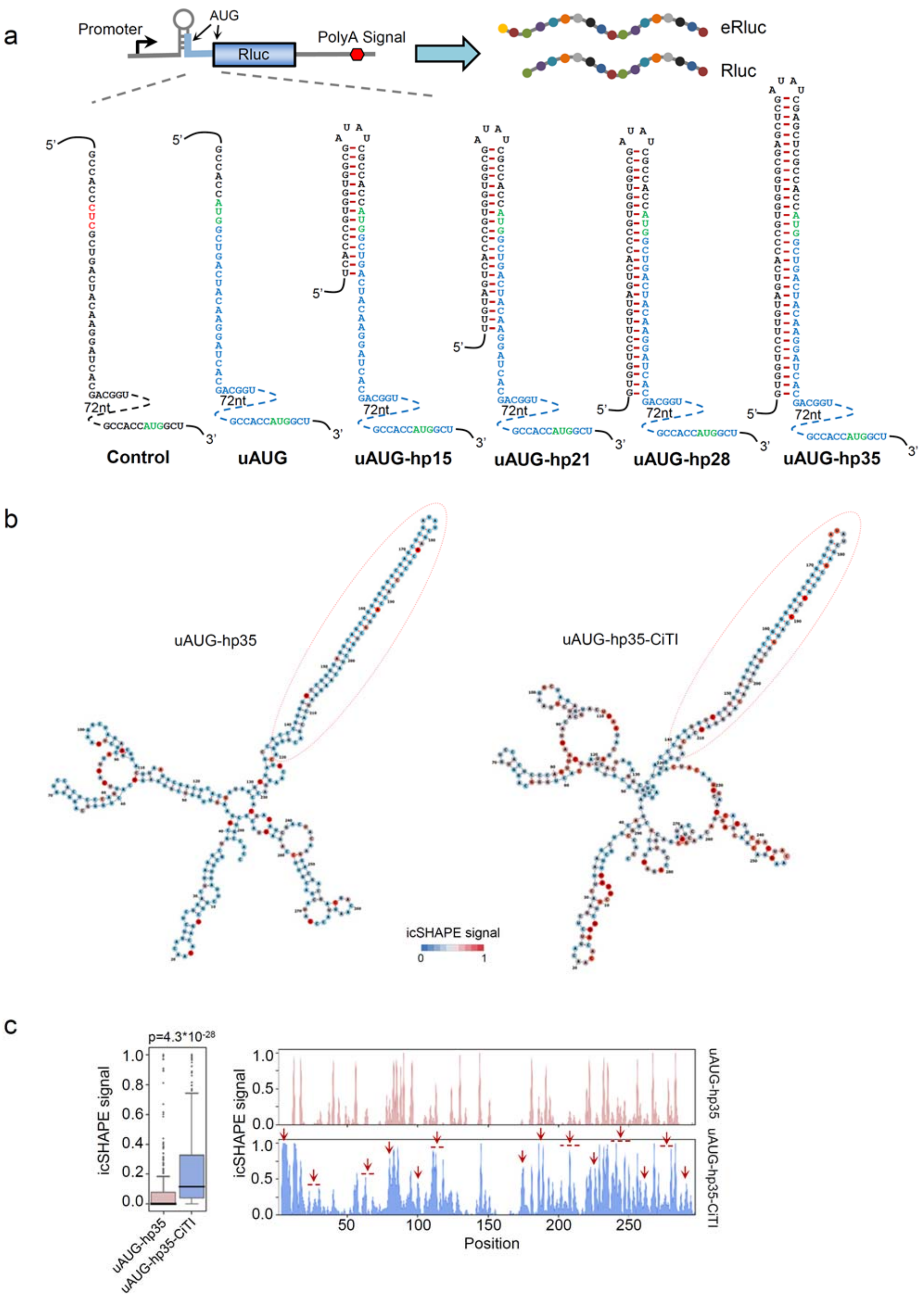


**Extended data figure 7. The 3′-CiTIs help to unwind 5′UTR structures. a,** Diagrams of the reporters with different structures in their 5′UTRs. The uAUG reporter contains an upstream start codon in the 5′UTR of Rluc ORF, which can generate an N-terminal extended coding region. In the control reporter, the upstream start codon was mutated to CUC; the additional reporters contain RNA hairpins of different lengths (15, 21, 28, 35 nt) in the 5′UTR of the uAUG reporter to mask the upstream start codon. The second green AUG represents the canonical start codon of Rluc. All reporters also contain a Fluc gene as an internal control. **b**, The RNA structures of the 5′UTR within uAUG-hp35 and uAUG-hp35-CiTI mRNAs. Full-length mRNAs were used to predict the RNA secondary structures based on the icSHAPE signals. The red circles indicate a high icSHAPE signal (i.e., a more "open" structure), and blue circles indicate a low icSHAPE signal (i.e. base-paired structures). **c**, *In vivo* icSHAPE signals of full-length 5′UTR regions in uAUG-hp35 and uAUG-hp35-CiTI mRNAs. The icSHAPE signals of the 5′UTR were box plotted (p values were calculated with unpaired two-sided Mann–Whitney U test). The regions with more opened structures after insertion of the CiTI are indicated by the red arrows.

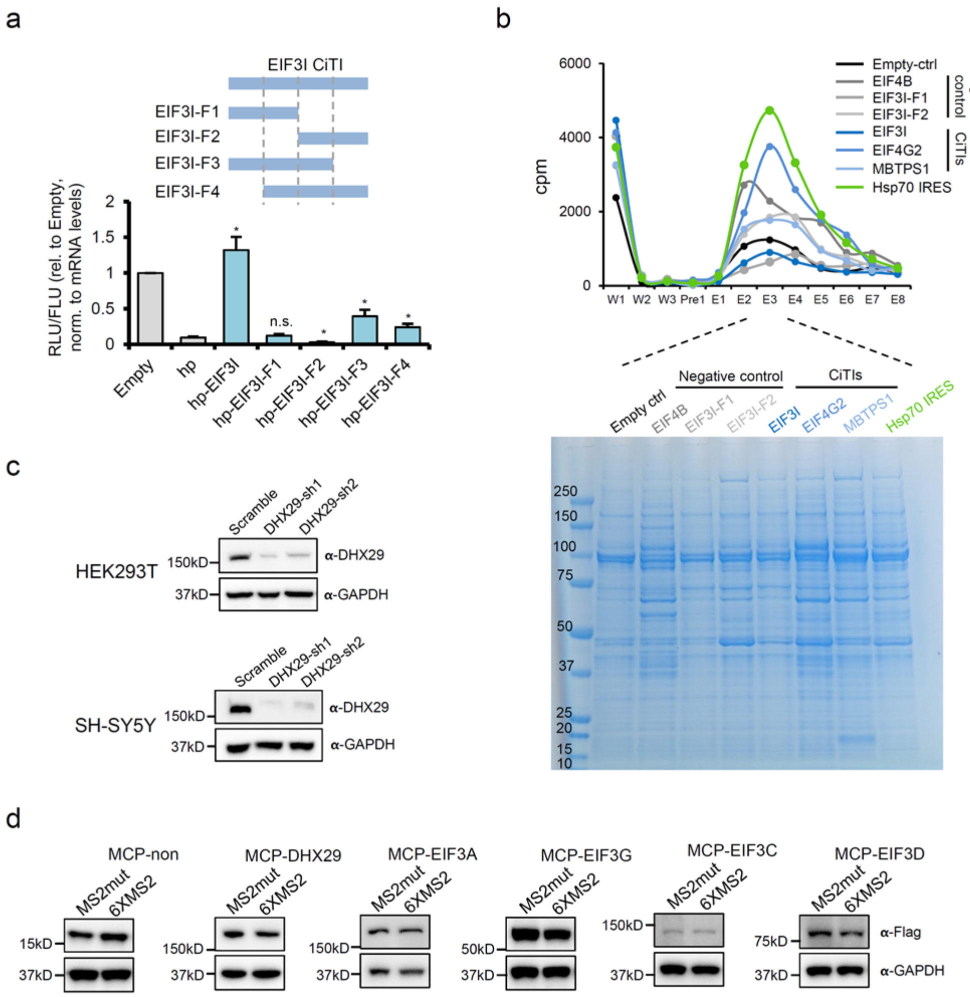


**Extended data figure 8. Identification of *trans*-factors binding to CiTIs. a**, The CiTI from eIF3I was split into different fragments, and inserted into the dual-luciferase reporters with a structured 5′UTR. Their activities were determined by translation efficiency, which was measured by the relative abundance of proteins (*via* luminescence assay) *vs.* RNAs (*via* qPCR) normalized to the control Fluc (n = 3, mean ± SD, * indicated p<0.05, n.s. indicated no significant difference by Student's t-test). **b**, RNAs containing MS2 aptamers were *in vitro* transcribed and used as "baits" to pull down *trans*-acting proteins (see details in Methods). Top, the relative abundance of the $^{32}$P-labeled RNAs in the wash (W1-3) and elutes (E1-E8) was determined by Cherenkov counting. Bottom, the proteins bound to each RNA bait were separated by SDS-PAGE and visualized by staining with Coomassie. Proteins in each lane were subsequently identified by mass spectrometry. **c**, The DHX29 gene was knocked down in HEK293 or SH-SY5Y cells using two individual shRNAs. DHX29 protein expression was judged by western blot in knockdown and scramble control cells. **d**, Measurement of the expression levels of different *trans*-factors fused with the MS2 coat protein. The fusion proteins were co-transfected with the translation reporters inserted with 6 copies of the wild type or mutated MS2 sequences, and the protein levels were determined by western blot.

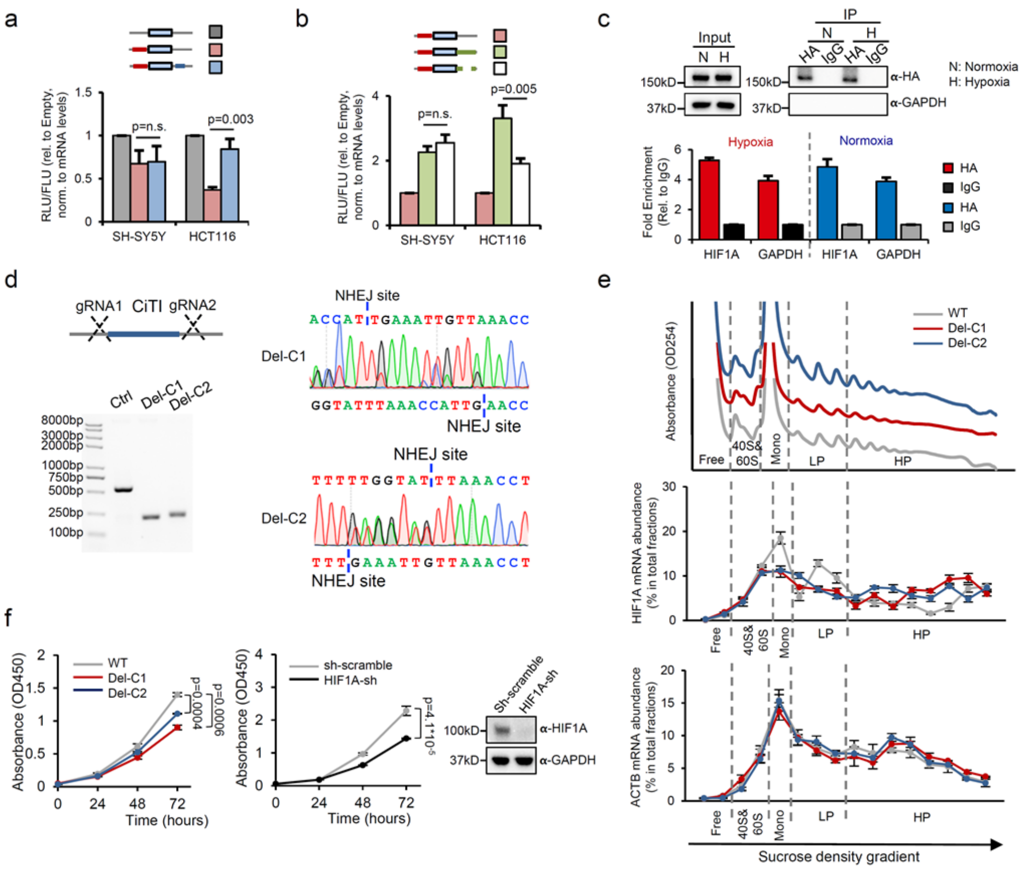


**Extended data figure 9. The function of the 3′-CiTI in HIF1A mRNAs under normal conditions. a**, The dual-luciferase reporters containing optionally the 5′UTR or 3′-CiTI of the HIF1A gene were transfected into SH-SY5Y or HCT116 cells under normal conditions (21% $O_2$). The translation efficiency of Rluc was determined by the relative abundance of proteins (*via* luminescence assay) *vs.* RNAs (*via* qPCR) normalized to the control Fluc (n = 3, mean ± SD, p value was calculated by Student's t-test). **b**, The full-length or CiTI-deleted 3′UTR of HIF1A was inserted into the Rluc gene containing the 5′UTR of HIF1A. The translation efficiency of Rluc under normal conditions in SH-SY5Y or HCT116 cells was determined as described in **panel a** (n = 3, mean ± SD, p value was calculated by Student′s t-test). **c**, The HA-tagged DHX29 expression vector was transfected into SH-SY5Y cells. The interactions of three endogenous mRNAs (HIF1A, GAPDH, and ACTB) and DHX29 were measured by RNA-IP and qPCR (n = 3, mean ± SD, n.s. indicated no significant difference by

Student's t-test). **d**, Validation of two individual clones with an HIF1A 3′-CiTI deletion. Left, agarose gel electrophoresis analysis of WT, Del-C1, and Del-C2′s PCR products. Right, sanger sequence chromatography of Del-C1 or Del-C2 PCR products. **e**, Two individual deletion clones (Del-C1 and Del-C2) and WT cells were collected and polysome profiling was performed under normal conditions. The total RNAs were extracted from each fraction and the abundance of HIF1A and ACTB mRNA in each fraction was further determined by RT-qPCR. Top, polysome profiling of WT and two individual deletion clones. Middle, the abundance of HIF1A mRNA in each fraction. Bottom, the abundance of ACTB mRNA in each fraction. **f**, Proliferation of WT or HIF1A 3′-CiTI-deleted SH-SY5Y cells under normal conditions. Cell proliferation was measured by a CCK8 assay. Left, cell proliferation of WT cells or two individual clones with an HIF1A 3′-CiTI deletion (n = 3, mean ± SD, p value was calculated by Student's t-test). Middle, the cell proliferation of HIF1A knockdown or scramble control cell line (n = 3, mean ± SD, p value was calculated by Student's t-test). Right, the protein expression of HIF1A in HIF1A knockdown or scramble cell lines at 1 h after hypoxia treatment.
